# AtTOC159 Receptors are Targeted to the Chloroplast Outer Membrane by a β-Signal and Galactolipid-Specific Transit Peptide-Like Sequence at the C-Terminus

**DOI:** 10.1101/2024.10.25.620236

**Authors:** Michael Fish, George Saudan, Simon D. X. Chuong, Masoud Jelokhani-Niaraki, Matthew D. Smith

## Abstract

Chloroplasts, the essential organelles responsible for photosynthesis, rely on the coordinated import of nuclear-encoded proteins for their biogenesis and function. TOC159 receptors are critical components of the translocon at the outer membrane of chloroplasts (TOC complex), responsible for recognizing and importing preproteins. However, the mechanisms by which TOC159 itself is targeted and integrated into the chloroplast outer envelope are only partially understood. In this study, we explore the structural aspects of the C-terminal membrane (M) domain of *Arabidopsis thaliana* TOC159, with a focus on its role in chloroplast outer envelope targeting and TOC complex integration. Structural predictions using AlphaFold and far-UV circular dichroism (CD) spectroscopy experiments suggested that the M domain forms an integral β-barrel that fuses with the β-barrel of TOC75, forming a hybrid channel at the chloroplast outer envelope. We identified a novel bi-partite targeting signal within the M domain, composed of a C-terminal β-signal with the consensus sequence G-Q-Φ-[ST]-Φ-[RK]-X-[SN]-[ST] and a transit peptide (TP)-like sequence. Using fluorescence microscopy and mutational analyses, we demonstrated that the β-signal is essential for chloroplast targeting, while the TP-like sequence enhances targeting efficiency. Furthermore, CD spectroscopy and Langmuir-Blodgett trough experiments revealed that the TP-like sequence preferentially interacts with galactolipids unique to the chloroplast outer membrane, particularly sulfoquinovosyl diacylglycerol (SQDG), facilitating chloroplast outer envelope targeting specificity. Overall, our findings provide a deeper understanding of TOC159 receptor targeting mechanisms, TOC complex biogenesis and highlight the broader significance of lipid-mediated targeting in intracellular protein trafficking. These results pave the way for future studies on the role of TP-like sequences in chloroplast outer membrane protein targeting and other novel targeting pathways more generally.

## 1. Introduction

Chloroplasts, the green engines within plant and algal cells, orchestrate the essential process of photosynthesis, converting light energy into chemical energy crucial to life (Jarvis and López-Juez, 2013). Photosynthetic eukaryotes inherited this ability through an ancient endosymbiotic event involving the acquisition of photosynthetic cyanobacteria. During their transition to the organelles we call plastids (including chloroplasts), most (thousands) of their genes were transferred to the nuclear genome (Day and Theg, 2018; Gould et al., 2008). The posttranslational targeting and import of chloroplast proteins are therefore critical processes to chloroplast biogenesis and function. Chloroplasts have evolved complex mechanisms to import newly synthesized “preproteins” from the cytosol. Typically, preproteins destined for chloroplasts contain a cleavable N-terminal transit peptide (TP) which interacts with a variety of cytosolic factors that recognize and direct preproteins to the chloroplast surface in an import-competent state (Bruce, 2000; Lee et al., 2008). Those cytosolic factors that have been characterized include heat shock protein (Hsp) 70 (Zhang and Glaser, 2002) and 90 (Qbadou et al., 2006), Hsp70/90-organising protein (Hop), and immunophilin FKBP73 (Fellerer et al., 2011). It has also been suggested that phosphorylated TPs associate with 14-3-3 proteins to form a guidance complex (Martin et al., 2006; May and Soll, 2000; Waegemann and Soll, 1996). As preproteins approach the chloroplast surface, translocation complexes at the outer (TOC) and inner (TIC) envelopes of the chloroplast mediate preprotein recognition and import (Kouranov and Schnell, 1997; Smith et al., 2004).

The core TOC complex is composed of the integral GTPase receptors, TOC159 and TOC33, and β-barrel import channel, TOC75 (Richardson et al., 2014). Numerous studies in *Arabidopsis thaliana* and *Pisum sativum* (pea) have documented the importance of each of these components in plant growth and development (Bauer et al., 2000; Bischof et al., 2011; Constan et al., 2004; Huang et al., 2011; Lee et al., 2003). TOC159 and TOC33 receptors possess homologous GTPase (G) domains that face the cytosol and work in tandem to recognize and bind chloroplast preproteins via their TPs (Koenig et al., 2008; Oreb et al., 2011; Sun et al., 2002). In *A. thaliana*, TOC159 (TOC132, TOC120, TOC90) and TOC33 (TOC34) homologs exist in structurally and functionally distinct TOC complexes (Ivanova et al., 2004), which exhibit different substrate preferences for photosynthetic and non-photosynthetic preproteins (Dutta et al., 2014; Grimmer et al., 2014; Infanger et al., 2011; Inoue et al., 2010; Kubis et al., 2004; Smith et al., 2004). TOC159 receptors also possess an N-terminal acidic (A) domain facing the cytosol that is intrinsically disordered (Richardson et al., 2009). Several studies have suggested that the A domain is responsible for the differential preprotein specificities of homologous TOC complexes (Agne and Kessler, 2010; Dutta et al., 2014). Once bound to the GTPase receptors, preproteins are directed to TOC75 for import (Agne and Kessler, 2010; Richardson et al., 2014). TOC75 is a member of the Omp85 superfamily, like those integral β-barrels in the outer membranes of gram-negative bacteria and mitochondria (Sommer et al., 2011). It contains several soluble N-terminal polypeptide transport-associated (POTRA) domains that extend into the intermembrane space, involved in chaperoning preproteins during translocation and as they engage the TIC complex (Chen et al., 2016; O’Neil et al., 2017).

Unlike chloroplast preproteins bound for import, most chloroplast outer envelope proteins (OEPs) lack N-terminal TPs. Instead, they rely on intrinsic targeting signals within their transmembrane domains (Fish et al., 2022). Signal- and tail-anchored proteins contain hydrophobic transmembrane ⍺-helices at their N- and C-termini, respectively. TOC33 and TOC34 receptors are examples of tail-anchored proteins (Teresinski et al., 2018). Specific cytosolic factors have been identified that direct these proteins to the chloroplast outer envelope for insertion by the TOC complex, including ankyrin repeat-containing protein 2 (AKR2) and small Hsp17.8 (sHsp17.8) (Kim et al., 2011; Teresinski et al., 2018). Chloroplast membranes are enriched in galactolipids such as monogalactosyl diacylglycerol (MGDG), digalactosyl diacylglycerol (DGDG) and sulfoquinovosyl diacylglycerol (SQDG), which are not found in other cellular membranes in plants (Schleiff et al., 2001). AKR2 interacts specifically with MGDG and phosphatidylglycerol (PG) head groups (Kim et al., 2014), highlighting the role of membrane lipid composition in chloroplast outer envelope targeting for these groups of proteins. The biogenesis of chloroplast outer membrane β-barrels is less understood, although one study has shown that OEP80 plays a key role in the insertion of this class of outer envelope protein (Gross et al., 2021), sharing analogous function with the β-barrel assembly machinery A (BamA) and sorting and assembly machinery 50 (sam50) proteins that facilitate β-barrel insertion in gram-negative bacteria and mitochondria, respectively (Höhr et al., 2018; Jores et al., 2016; Kutik et al., 2008; Robert et al., 2006). To date, only two chloroplast outer envelope proteins (TOC75 and OEP80) have been shown to use cleavable N-terminal TPs, despite their integral β-barrel structure (Day et al., 2019; Gross et al., 2020).

While the role of TOC159 receptors in preprotein recognition is well established, the mechanisms governing its own targeting and integration into the chloroplast outer envelope have been mostly limited to the role of an interaction between cognate G-domains of TOC159 and TOC33 and the GTP-status of the TOC159 G-domain (Bauer et al., 2002; Smith et al., 2002; Wallas et al., 2003). The relative lack of understanding about the mechanism of TOC159 targeting was, until very recently, due to a poor understanding of the C-terminal membrane (M) domain. With no predictable transmembrane structures, but resistance to proteolysis and alkaline extraction from the chloroplast outer envelope (Bauer et al., 2000; Bölter et al., 1998), it was thought to form an unconventional membrane anchor (Lung et al., 2014). Recent predictions by AlphaFold and cryo-EM structures of the TOC complex from the green algae, C*hlamydomonas reinhardtii* shine an entirely new light on the structure of the TOC159 M domain, as an integral β-barrel which fuses with the β-barrel of TOC75 to form a hybrid channel structure reminiscent of the substrate-bound BamA complex in the cyanobacterial endosymbiont (Jin et al., 2022; Liu et al., 2023). This revolutionary discovery shifts our focus from an impartial or purely structural membrane anchor to an active pore, which may play a larger role in protein import than earlier thought.

Previously, we characterized three subdomains within the M domain of TOC159 receptors: the M1 subdomain is largely disordered, containing a predicted coiled-coil; the M2 or Mβ subdomain is composed of the first 12 β-strands that are now predicted to form the β-barrel; and the M3 subdomain includes the last 2 β-strands of the predicted β-barrel followed by a C-terminal sequence that shares similar physicochemical characteristics with the N-terminal TPs of chloroplast preproteins (Lung et al., 2014). We demonstrated that the M3 subdomain from *Bienertia sinuspersici* and *A. thaliana* was necessary, but not sufficient for localization to the chloroplast outer envelope. As more of the β-strands from Mβ were included, localization of enhanced green fluorescent protein (eGFP) fusion proteins to the chloroplast outer envelope improved, demonstrating a reversible association with the chloroplast outer envelope followed by irreversible anchorage (Lung and Chuong, 2012; Lung et al., 2014).

Building on these findings and considering the new structural evidence, the current study aimed to establish that TOC159 receptors are anchored in the chloroplast outer membrane by a β-barrel in plants and analyze the role of individual elements within the M3 subdomain, as well as the mechanisms they employ in reversible chloroplast outer envelope association and membrane integration. We hypothesized that the β-hairpin structure or ultimate β- strand within the M3 subdomain could constitute a β-signal responsible for targeting like those of β-barrels in the outer membranes of mitochondria and gram-negative bacteria. Further, we sought to identify the role of galactolipids in the reversible interaction of the TP-like sequence. To achieve this, we used fluorescence microscopy to observe the subcellular localization of various fluorescent fusion proteins transiently expressed in *Allium cepa* (onion) epidermal cells and *A. thaliana* mesophyll cell protoplasts to understand the role of the β-hairpin, ultimate β-strand and TP-like sequence individually in chloroplast outer envelope targeting. Additionally, we used far-UV circular dichroism (CD) spectroscopy and Langmuir-Blodgett trough experiments to evaluate the interactions between the TP-like sequence and galactolipids. Together, our findings showed that the C-terminus of AtTOC159 receptors from *A. thaliana* contain a conserved bi-partite targeting signal, composed of a β- signal and TP-like elements that preferentially interact with galactolipids to direct the protein to the chloroplast outer envelope.

## 2. Materials and Methods

### 2.1 Bioinformatic Analyses

The structure of the AtTOC complex was predicted using ColabFold v1.5.5: AlphaFold2 using MMseqs2 (Evans et al., 2021; Mirdita et al., 2022, 2019, 2017; Mitchell et al., 2019). The predicted structures of AtTOC159, AtTOC132, AtTOC120 and AtTOC90 were retrieved from the AlphaFold Protein Structure Database (Jumper et al., 2021; Varadi et al., 2024). Protein BLAST was used to generate a list of all available homologous TOC159 protein sequences across plant species (Altschul et al., 1990). Putative, uncharacterized and hypothetical proteins were removed, and 1104 sequences were aligned using the ClustalW algorithm (Madeira et al., 2024). This alignment was converted to a sequence logo using WebLogo 3.7.12 (Crooks et al., 2004; Schneider and Stephens, 1990). Helical wheel analyses were performed using HeliQuest to determine hydrophobicity (H), hydrophobic moment (µH), and net charge (z) (Gautier et al., 2008).

### 2.2 Construct Generation

*AtTOC159* (AT4G02510) and *AtTOC132* (AT2G16640) sequences were synthesized and sub-cloned into the pSAT6-35S:eGFP-C1 vector (Lung and Chuong, 2012) by BioBasic Inc. (Canada) to generate eGFP fusion constructs. The DsRed2 construct used in biolistic bombardment of onion epidermal cells was generated previously by excising the TP sequence of ferredoxin from a previous construct and sub-cloning it into the pSAT6- 35S:DsRed2-N1 vector at the 5′ end of the DsRed2-encoding sequence (Chung et al., 2005; Lung et al., 2011). *AtTOC75* (AT3G46740), *AtTOC159* and *AtTOC132* sequences encoding only the β-barrel region containing a poly-histidine tag at their C-termini were synthesized and sub-cloned into the pET28a(+) vector by BioBasic Inc. (Canada) to generate constructs for recombinant protein expression in *Escherichia coli*. Sequences used in the generation of constructs are available in **Tables S1 and S2** of the Supplemental Materials. All constructs were verified using DNA sequencing.

### 2.3 Expression, Purification & Refolding of AtTOC75, AtTOC159 and AtTOC132 β-Barrels

Recombinant, poly-histidine tagged AtTOC75, AtTOC159 and AtTOC132 β-barrels were overexpressed in *E. coli* BL21 (DE3) LOBSTR cells by IPTG induction and purified from inclusion bodies using immobilized metal affinity chromatography with HisPur^TM^ Ni-NTA resin (Thermo Scientific, 88221). Before elution, proteins were refolded in buffer (20 mM Tris-HCl, pH 8.0, 50 mM NaCl, 15 mM imidazole) containing 0.1% Triton X-100 detergent by on-column, stepwise buffer exchange as described by Rogl et al. (1998). Proteins were eluted with 400 mM imidazole and desalted into CD1 Buffer (20 mM Tris-HCl, pH 8.0, 50 mM NaCl, 0.1% Triton X-100) using Econo-Pac 10DG Desalting Columns (Bio-Rad, 7322010). Fractions were analyzed by Bradford protein assay and SDS-PAGE (4% stacking and 12% resolving gels). Gels were stained with Coomassie Brilliant Blue and imaged using a ChemiDoc MP Imaging System (Bio-Rad, Canada).

### 2.4 Biolistic Bombardment of *Allium cepa* (Onion) Epidermal Cells

Bulbs of *A. cepa* (onion) were purchased from local grocery stores and stored at 4 ℃. The plasmid DNA (5 µg) of eGFP and DsRed2 fusion constructs were coated on tungsten particles (1.1 µm in diameter; Bio-Rad, 165-2267) in a suspension containing 16 mM spermidine and 0.1 M CaCl_2_. The plasmid DNA-coated tungsten particles were dried on microcarrier discs (Bio-Rad, 165-2335) and bombarded into the adaxial surface of onion bulb sections (1 cm^2^) from a distance of 12 cm at a pressure of 1350 psi using a Biolistic PDS- 1000/He particle delivery system (Bio-Rad, 165-2257). Bombarded onion samples were incubated in petri dishes on moist filter paper at room temperature in the dark for 14 hours and observed using epifluorescence microscopy.

### 2.5 Epifluorescence Microscopy

Epifluorescence microscopy of *A. cepa* (onion) epidermal cells was carried out using a Zeiss Axio Imager D1 microscope equipped with a Zeiss AxioCam MRm Camera (Carl Zeiss Inc., Germany). eGFP and DsRed2 signals were acquired using excitation wavelengths of 470 nm and 550 nm, respectively and emission wavelengths of 525 nm and 570 nm, respectively. Representative images (n = 3) were selected and rendered using AxioVision 4.7 software. The Pearson’s correlation (R) between eGFP and DsRed2 channels in merged images was determined using “Coloc 2” analysis in Image J (Fiji) software (Schindelin et al., 2012). Mean correlation coefficients (n = 3) ± the standard error of the mean (SEM) were analyzed using one-way ANOVA and the Tukey multiple comparisons test with the BioRender R 4.2.2 Graph integration.

### 2.6 Plant Materials & Growth Conditions

*A. thaliana* seeds (ecotype Columbia-0) were stratified in a solution of 0.1% agarose at 4 ℃ in the dark for 48 hours before being sown on the surface of pots containing Sunshine Mix #4 soil (Sungro Horticulture). Pots were covered with transparent lids until cotyledons were observed, after which the covers were removed, and the plants were watered from the bottom regularly. Plants were maintained in a growth chamber with a light/dark cycle of 12/12 hours at 22 ℃ with a photon flux density of ca. 100 µmol m^-2^ s^-1^. True leaves from 3- to 4-week-old plants were used for protoplast isolation and transfection.

### 2.7 *Arabidopsis thaliana* Protoplast Isolation & Transfection

Mesophyll protoplasts were isolated from *A. thaliana* using the tape-leaf sandwich method (Wu et al., 2009). True leaves from 3- to 4-week-old plants were fixed to Scotch Masking Tape with their lower epidermis facing upwards. Strips of Scotch Magic Tape were placed over the leaves and used to remove the lower epidermis. As described previously by Yoo et al. (2007), the resulting leaves with exposed mesophyll cells were incubated in enzyme solution (0.4 M mannitol, 20 mM MES-KOH, pH 5.7, 20 mM KCl, 10 mM CaCl_2_, 0.1% [w/v] bovine serum albumin, 1.0% [w/v] cellulase Onozuka R10 and 0.25% [w/v] macerozyme R10 enzymes from Yakult Pharmaceutical, Japan) with shaking at room temperature in the dark for 1 hr. The isolated protoplasts were washed twice with equal volume of W5 solution (2 mM MES-KOH, pH 5.7, 154 mM NaCl, 125 mM CaCl_2_ and 5 mM KCl) and the protoplasts were pelleted by centrifugation at 100 g for 3 min. The protoplast pellets were resuspended in CS-Sucrose solution (0.4 M Sucrose, 20 mM MES-KOH, pH 5.7, 20 mM KCl) and healthy protoplasts were isolated by centrifugation at 100 g for 3 min. The healthy protoplasts in the floating layer were removed, diluted in 1 mL of W5 solution and allowed to settle for 30 min in the dark at 4℃. The settled protoplasts were resuspended in Mg-Man buffer (0.4 M mannitol, 4 mM MES-KOH, pH 5.7, 15 mM MgCl_2_) at a density of 200,000 protoplasts mL^−1^. Protoplast viability was determined using fluorescein diacetate staining under epi-fluorescence microscopy. Protoplasts were transfected based on a modified protocol from Yoo et al. (2007). Approximately 200,000 protoplasts were mixed with 50 μg of plasmid DNA and 1 mL of polyethylene glycol solution (40% [w/v] PEG4000 from Sigma-Aldrich, 0.4 M sucrose, 0.1 M CaCl_2_). After incubation at room temperature for 15 min, the transfected protoplasts were mixed with 4 mL of W5 solution, pelleted at 100 g for 3 min, resuspended in 4 mL of WI solution (0.5 M mannitol, 4 mM MES-KOH, pH 5.7, 20 mM KCl) and cultured overnight at 23◦C with a light intensity of 30 μmol m^−2^ s^−1^.

### 2.8 Confocal Laser Scanning Fluorescence Microscopy

Transformed *A. thaliana* protoplasts were examined in a Thermo Scientific™ Nunc™ Lab-Tek™ II Chamber Slide™ (Thermo Scientific, 154453) using an inverted Olympus FluoView 1000 confocal laser-scanning microscope (Olympus Co., Japan). eGFP and chlorophyll autofluorescence signals were acquired simultaneously using excitation wavelengths of 488 nm and 543 nm, respectively and emission bandwidths of 505-525 nm and 655-755 nm, respectively. Representative images (n = 3) were selected from z-stacks of optical sections (1.0 µm optical slice thickness) and rendered using the Olympus Fluoview imaging software (FV10-ASW 1.7).

### 2.9 (Proteo)Liposome Preparation

Recombinant AtTOC75, AtTOC159 and AtTOC132 β-barrels were reconstituted into palmitoyl oleoyl phosphatidylcholine (POPC) liposomes using the detergent-mediated reconstitution method as described previously (Tabefam et al., 2024). In brief, rehydrated lipids were solubilized in C_8_E_4_ detergent and combined with protein in CD1 Buffer. Spontaneous formation of proteoliposomes occurred after incubation with SM-2 Biobeads (Bio-Rad, 1523920) to remove the detergent. The AtTOC159M3 peptide was synthesized by BioBasic Inc. (Canada). Liposomes used in peptide-lipid interaction studies were prepared by the extrusion method (MacDonald et al., 1991) in CD2 Buffer (20 mM Tris-HCl, pH 8.0, 50 mM NaF). Rehydrated lipids were passed through a mini-extruder device (Avanti Polar Lipids Inc., USA) using a 0.1 µm filter to produce large unilamellar vesicles of homogenous size. Liposomes used in these studies were composed of POPC, POPC:MGDG (19:1), POPC:DGDG (19:1), POPC:SQDG (19:1), and POPC:MGDG:DGDG:SQDG (5:2.5:5:1). The composition of POPC:MGDG:DGDG:SQDG liposomes was chosen to mimic the chloroplast outer membrane (Schleiff et al., 2001). All lipids were purchased from Avanti Polar Lipids, Inc. (USA).

### 2.10 Far-UV Circular Dichroism (CD) Spectroscopy

CD spectra were obtained from 190-260 nm at 1 nm resolution using an AVIV 215 spectropolarimeter (AVIV Instruments Inc., USA). Protein samples (∼8 µM in detergent and ∼1 µM in proteoliposomes) were measured in and corrected against CD1 Buffer. Peptide samples (10 µM) were measured in and corrected against CD2 Buffer. All samples were measured at 22 °C in a 0.1 cm path-length quartz cuvette. The reported spectra are converted to mean residue ellipticity (MRE) [*θ*] in degrees cm^2^ dmol^-1^ and represent an average of 9 scans (n = 3).

### 2.11 Langmuir-Blodgett Trough Experiments

In a Langmuir-Blodgett trough, 10 µg of total lipid – POPC, MGDG, DGDG, SQDG, or a mixture of POPC:MGDG:DGDG:SQDG (5:2.5:5:1) – dissolved in chloroform was spread onto the subphase (10 mM Tris-HCl pH 7.4, 150 mM NaCl). After allowing the chloroform to evaporate for 20 minutes, the trough barriers were compressed to an initial surface pressure (πi) of 0 mN/m. Then, 10 µM of synthetic AtTOC159 TP-like peptide was added to the subphase. Peptide was injected over a range of πi, and changes in surface pressure (Δπ) were monitored every 0.25 seconds for ∼4 minutes. Δπ was plotted against πi, and linear regression was used to extrapolate the x-intercept, indicating the maximum insertion pressure of the peptide in each lipid environment. The mean maximum insertion pressures (n = 3) ± standard error of the mean (SEM) were analyzed using one-way ANOVA and the Tukey multiple comparisons test with the BioRender R 4.2.2 Graph integration.

## 3. Results

### 3.1 AtTOC159 Receptors are Anchored in the Chloroplast Outer Envelope by a β-Barrel

To determine whether TOC159 receptors in plants adopt β-barrel membrane anchors as they do in algae, ColabFold was used to predict the structure of the AtTOC complex. As seen in **Figure 1A** and **Figure 1B**, the M domain of AtTOC159 was predicted by AlphaFold to form a β-barrel membrane anchor that is fused to the β-barrel of AtTOC75, just like in the cryo-EM structure of the CrTOC complex in algae (Jin et al., 2022; Liu et al., 2023). The M1 and M3 subdomains were also predicted to form structurally distinct regions as previously predicted (Lung et al., 2014) – the M1 subdomain contained a coiled-coil and the M3 subdomain contained an ⍺-helix. Additionally, the M domains of AtTOC132, AtTOC120 and AtTOC90 were predicted to adopt the same structure with root mean square deviations of 0.866, 0.979 and 1.064 angstroms relative to the M domain of AtTOC159, respectively (**Figure 1C**). The β- barrel region of AtTOC75 and the predicted β-barrel regions of AtTOC159 and AtTOC132 were recombinantly expressed in *Escherichia coli* and purified using affinity chromatography (**Figure S1**). CD spectroscopy was used to compare the secondary structure of each in Triton X-100 detergent and when reconstituted in POPC liposomes. The CD spectra (**Figure 1 D-F**) showed that both the predicted β-barrel regions of AtTOC159 and AtTOC132 have similar structures to the β-barrel region of AtTOC75 in both environments, with minima around 215 nm, characteristic of β-barrels (Greenfield, 2006; Miles et al., 2021).

**Figure 1.**
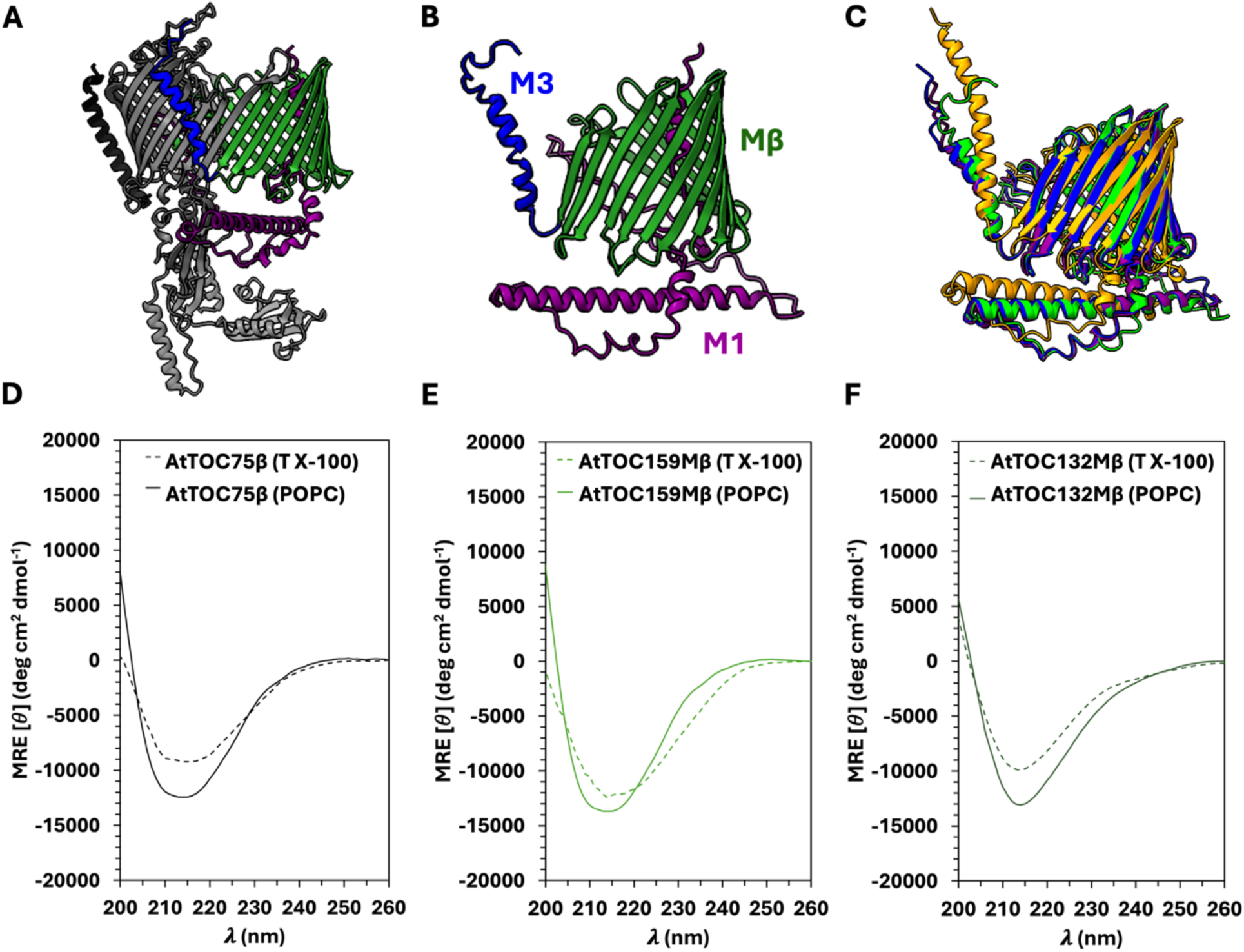
Structural Characterization of the AtTOC159 Receptor Membrane Anchor. (**A**) Structural prediction of the TOC complex from *Arabidopsis thaliana* using ColabFold v1.5.5: AlphaFold2 with MMseqs2 (Evans et al., 2021; Mirdita et al., 2022, 2019, 2017; Mitchell et al., 2019). AtTOC34 is depicted in black, AtTOC75 is depicted in grey and the AtTOC159 membrane (M) domain is coloured by subdomain (M1 in purple, Mβ in green and M3 in blue). (**B**) AlphaFold predicted structure of the AtTOC159 M domain (coloured as in **A**) and (**C**) superimposed AtTOC159 homologous (AtTOC159, AtTOC132, AtTOC120 and AtTOC90) M domains (Jumper et al., 2021; Varadi et al., 2024). Protein structures were generated and aligned using UCSF Chimera. (**D-F**) Far-UV CD spectra of the recombinantly expressed and purified β-barrel regions of AtTOC75 (**D**), AtTOC159 (**E**) and AtTOC132 (**F**) in 0.1% Triton X-100 detergent (dashed lines) and reconstituted in POPC liposomes (solid lines). All spectra, considering the structural predictions, confirm a β-barrel structure. The protein concentration in (**D-F**) was 8 µM in detergent and 1 µM in proteoliposomes. All spectra are an average of 9 scans (n = 3) and were collected at 22 ℃ in a quartz cuvette with a 0.1 cm pathlength. MRE is the mean residue ellipticity (*θ*) in degrees cm^2^ dmol^-1^.

In previous studies, we defined the M3 subdomain according to its chloroplast outer envelope targeting capabilities and the observation that this region shares similar physicochemical characteristics with chloroplast TPs (Lung and Chuong, 2012; Lung et al., 2014). It is now clear that the β-strands within M3 represent the last two β-strands of the β- barrel and should be included in our definition of the Mβ subdomain instead, redefining the M3 subdomain to include the C-terminal ⍺-helix alone. The targeting of β-barrel outer membrane proteins of the mitochondria and gram-negative bacteria is well characterized, dependent on the C-terminal β-strand of the β-barrel known as a β-signal (Höhr et al., 2018; Jores et al., 2016; Kutik et al., 2008; Robert et al., 2006). Multiple sequence alignment of TOC159 homologs across plant species revealed that the last β-strand of the β-barrel is highly conserved, with the consensus sequence G-Q-Φ-[ST]-Φ-[RK]-X-[SN]-[ST] (**Figure 2A**), where X represents any amino acid and Φ represents a hydrophobic residue. Although the M3 subdomain is less conserved, the predicted ⍺-helix and physicochemical characteristics shared with chloroplast TPs imparted by the amino acid residues at each location are well conserved. Further, helical wheel projections of the predicted ⍺-helices within the M3 subdomain of the AtTOC159 receptors demonstrated a conserved amphipathic nature, with a hydrophobic face and a positively charged hydrophilic face (**Figure 2B**). The hydrophobicity and hydrophobic moment were determined using HeliQuest for each ⍺-helix: AtTOC159 had a hydrophobicity of 0.839 and a hydrophobic moment of 0.291, AtTOC132 had a hydrophobicity of 0.838 and a hydrophobic moment of 0.330, AtTOC120 had a hydrophobicity of 0.848 and a hydrophobic moment of 0.308, and AtTOC90 had a hydrophobicity of 0.578 and a hydrophobic moment of 0.233.

**Figure 2.**
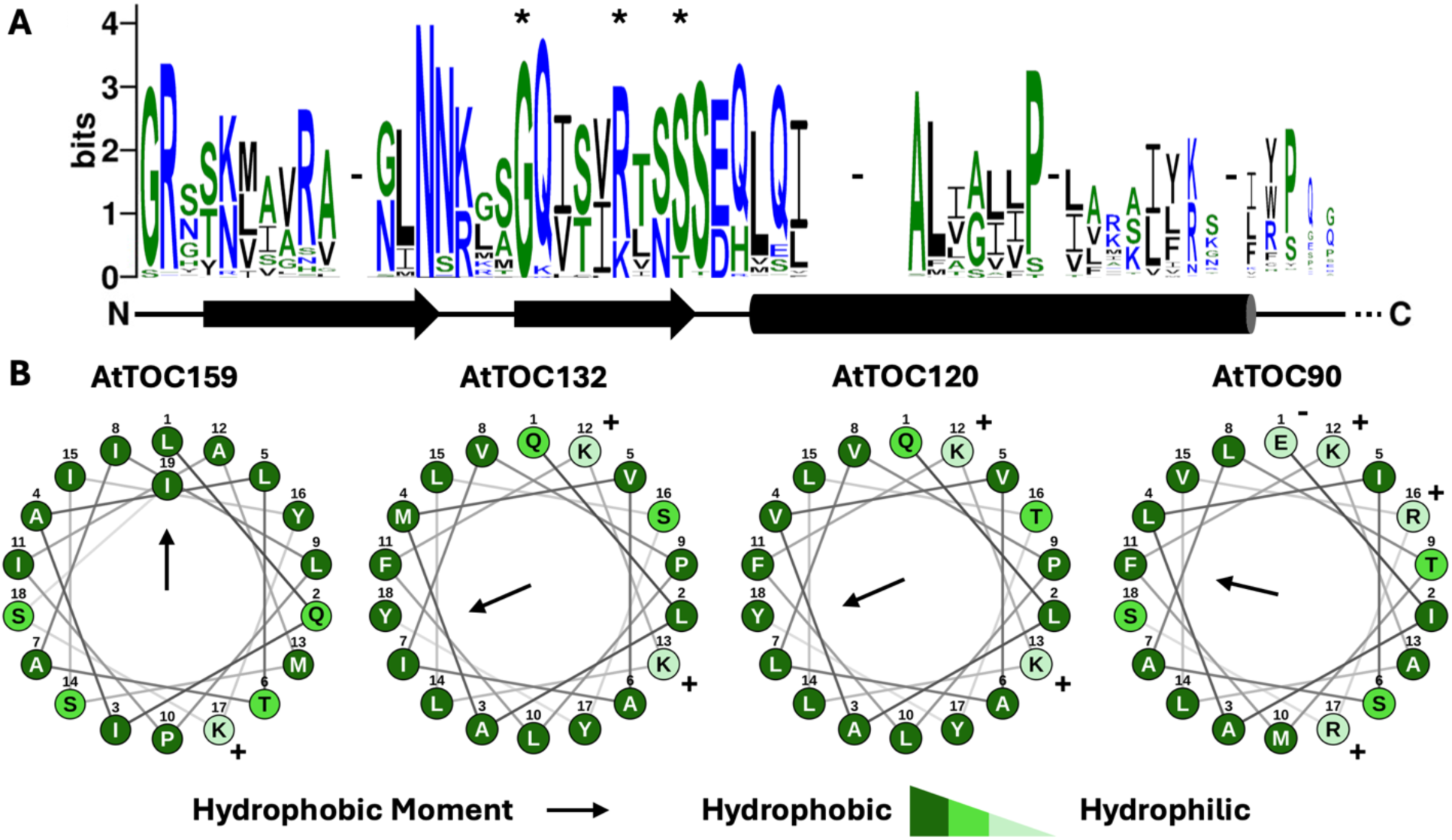
Bioinformatic Analysis of the C-Terminal Region of TOC159 Receptor Homologs in Plants. (**A**) WebLogo 3.7.12 output of the aligned C-terminal sequences of TOC159 homologs (Crooks et al., 2004; Schneider and Stephens, 1990). 1104 sequences were acquired using BlastP (Altschul et al., 1990) and aligned using the ClustalW algorithm (Madeira et al., 2024). The height of each amino acid residue represents the frequency in the aligned sequences. Gaps in the sequence alignment are highlighted by a dash (-). Blue residues are hydrophilic, green residues are neutral and black residues are hydrophobic. The secondary structure as predicted by AlphaFold in the AtTOC159 structure is depicted below the output, illustrating the high level of conservation in the last β-strand of the predicted β-barrel. The 3 residues chosen for mutation to alanine in this study are marked with an asterisk (*). (**B**) Helical wheel analysis of the predicted C-terminal ⍺-helices of AtTOC159 homologs. The sequences of each ⍺-helix as predicted by AlphaFold were analyzed using HeliQuest (Gautier et al., 2008) to determine the helical wheel projection and hydrophobic moment, illustrating a conserved amphipathic nature with a hydrophobic face and positively charged hydrophilic face. Residues in the helical wheel projections are coloured with charged residues in light green, polar residues in green and hydrophobic residues in dark green.

### 3.2 AtTOC159 Receptors Contain a Novel Bi-Partite Targeting Signal at the C-Terminus

Considering our newly proposed definitions of the Mβ and M3 subdomains, we used eGFP fusion constructs (**Figure 3A**) to evaluate the plastid targeting capabilities of each subdomain in *A. cepa* and the chloroplast outer envelope targeting capabilities of each subdomain in *A. thaliana* protoplasts. The biolistic bombardment of *A. cepa* epidermal cells provides a high-throughput analysis of plastid colocalization of eGFP fusion constructs and the DsRed plastid marker using epi-fluorescence microscopy (**Figure 3B**). Further, the plastid colocalization of eGFP fusion constructs and the plastid marker were quantified using “Coloc 2” analysis in Image J (Fiji) software (**Figure 5**). As seen in **Figure 3B** and **Figure 5A**, three significant levels of targeting capability were observed: good, reduced and poor. When the M1 subdomain is present with other subdomains (AtTOC159M, AtTOC159Mβ and AtTOC159MΔM3), targeting capability was reduced, which suggests that the M1 subdomain interferes with targeting in the context of these experiments. AtTOC159M1 on its own demonstrated poor targeting capability, further confirming that the M1 subdomain is not involved in targeting. On the other hand, AtTOC159MΔM1, which contains both the Mβ and M3 subdomains, showed good targeting capability. Unexpectedly, the observation that AtTOC159MΔM3 showed some targeting capability and AtTOC159M3 showed poor targeting capability suggests that the M3 subdomain is not essential for targeting but contributes to targeting efficiency. Further, the observation that AtTOC159MΔMβ showed poor targeting capability and AtTOC159Mβ showed some targeting capability suggests that the Mβ subdomain contains essential targeting elements.

**Figure 3.**
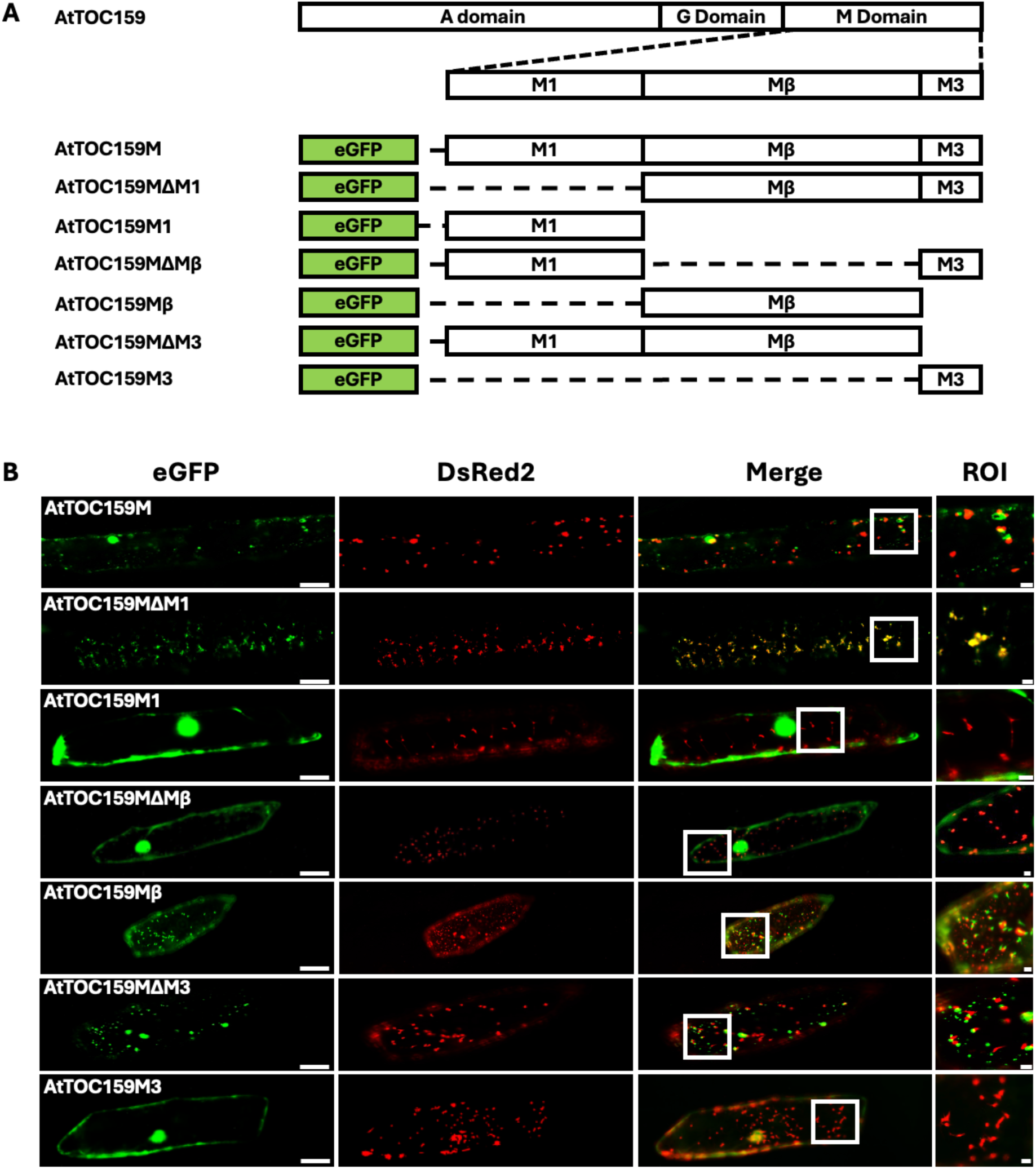
Subcellular Localization of Transiently Expressed eGFP-AtTOC159 Fusion Constructs Containing the M Domain, Subdomains and Subdomain Truncations in *Allium cepa* (onion) Epidermal Cells. (**A**) Schematic Representation of eGFP-AtTOC159 Fusion Constructs. AtTOC159 constructs were fused to the C-terminus of eGFP in the pSAT6-35S:eGFP-C1 vector to be transiently expressed using the 35S promoter. (**B**) The epidermal cells of onion bulbs were co-transfected with eGFP-AtTOC159 fusion constructs and a DsRed2-Ferredoxin transit peptide fusion construct using biolistic bombardment. Representative images for each co-bombardment include eGFP (green), DsRed2 (red) and the merged channels. Colocalization of the eGFP and DsRed2 fusion proteins to plastids produces a yellow signal. The region of interest (ROI) is representative of the areas in the merged image used to analyze colocalization (Figure 5). Scale bars = 50 µm (eGFP) and 5 µm (ROI).

Additional eGFP fusion constructs (**Figure 4A**) were made to probe the role of the predicted β-signal by truncation or mutagenesis. Ultimately, truncations of the β-strand of the β-barrel including upstream (AtTOC159MΔβH) and downstream (AtTOC159MΔCS) residues abolished targeting. Additionally, mutation of 3 conserved residues in the β-signal consensus sequence to alanine (AtTOC159M3A) also abolished targeting. This established the β-signal as an essential targeting signal. Furthermore, AtTOC159MΔβH and AtTOC159MΔCS were not capable of targeting, suggesting that although essential, the β- signal is not sufficient for targeting. These results were also observed for AtTOC132 eGFP fusion constructs (**Figure S2**).

**Figure 4.**
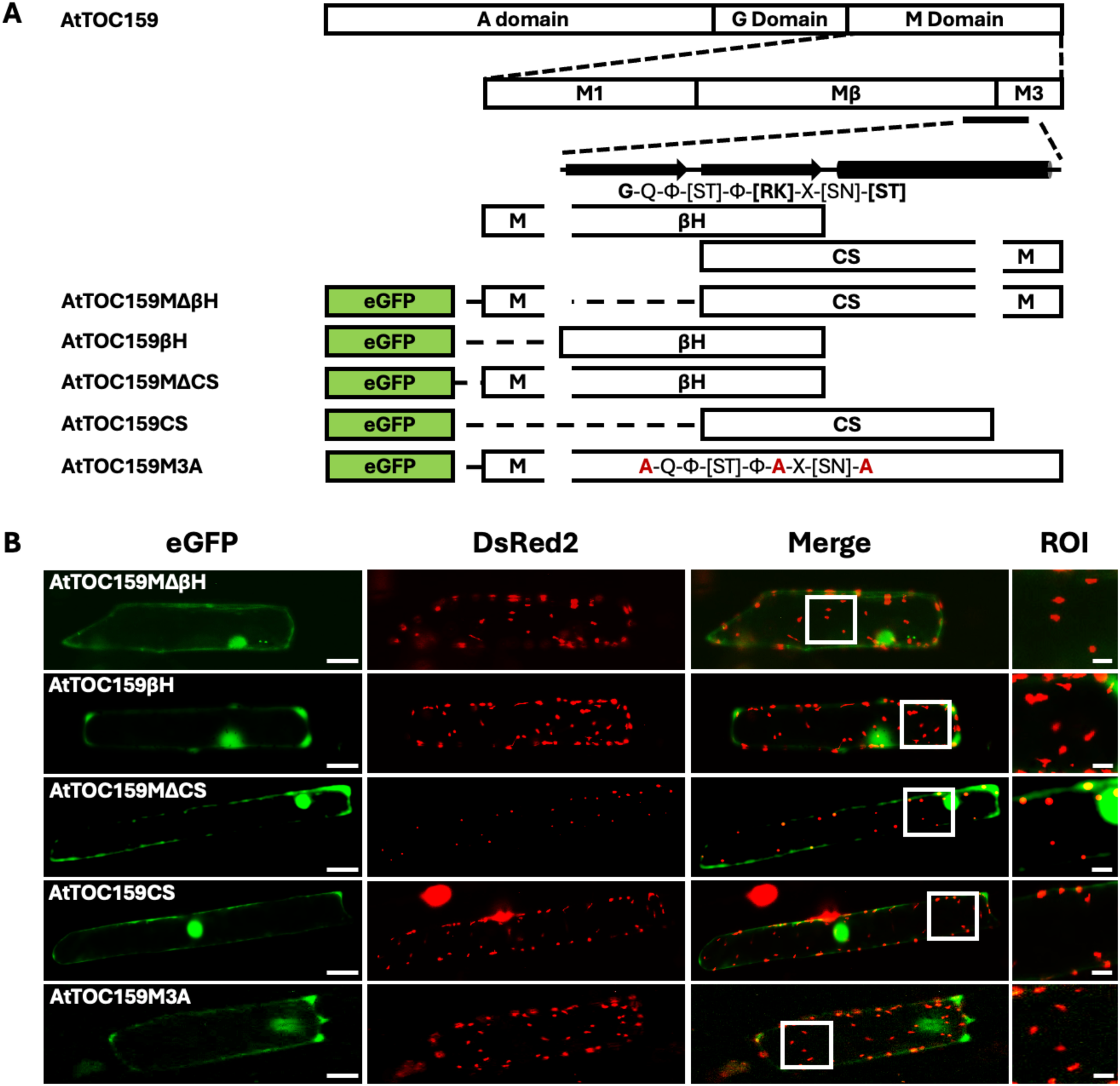
Subcellular Localization of Transiently Expressed eGFP-AtTOC159 Fusion Constructs Containing C-Terminal Regions, Truncations and Alanine Mutations in *Allium cepa* (onion) Epidermal Cells. (**A**) Schematic Representation of eGFP-AtTOC159 Fusion Constructs. AtTOC159 constructs were fused to the C-terminus of eGFP in the pSAT6-35S:eGFP-C1 vector to be transiently expressed using the 35S promoter. The conserved motif corresponding to the last β-strand of the predicted β-barrel and corresponding alanine mutations in the eGFP-AtTOC159M3A fusion construct are highlighted. (**B**) The epidermal cells of onion bulbs were co-transfected with eGFP-AtTOC159 fusion constructs and a DsRed2-Ferredoxin transit peptide fusion construct using biolistic bombardment. Representative images for each co-bombardment include eGFP (green), DsRed2 (red) and the merged channels. Colocalization of the eGFP and DsRed2 fusion proteins to plastids produces a yellow signal. The region of interest (ROI) is representative of the areas in the merged image used to analyze colocalization (Figure 5). Scale bars = 50 µm (eGFP) and 5 µm (ROI).

**Figure 5.**
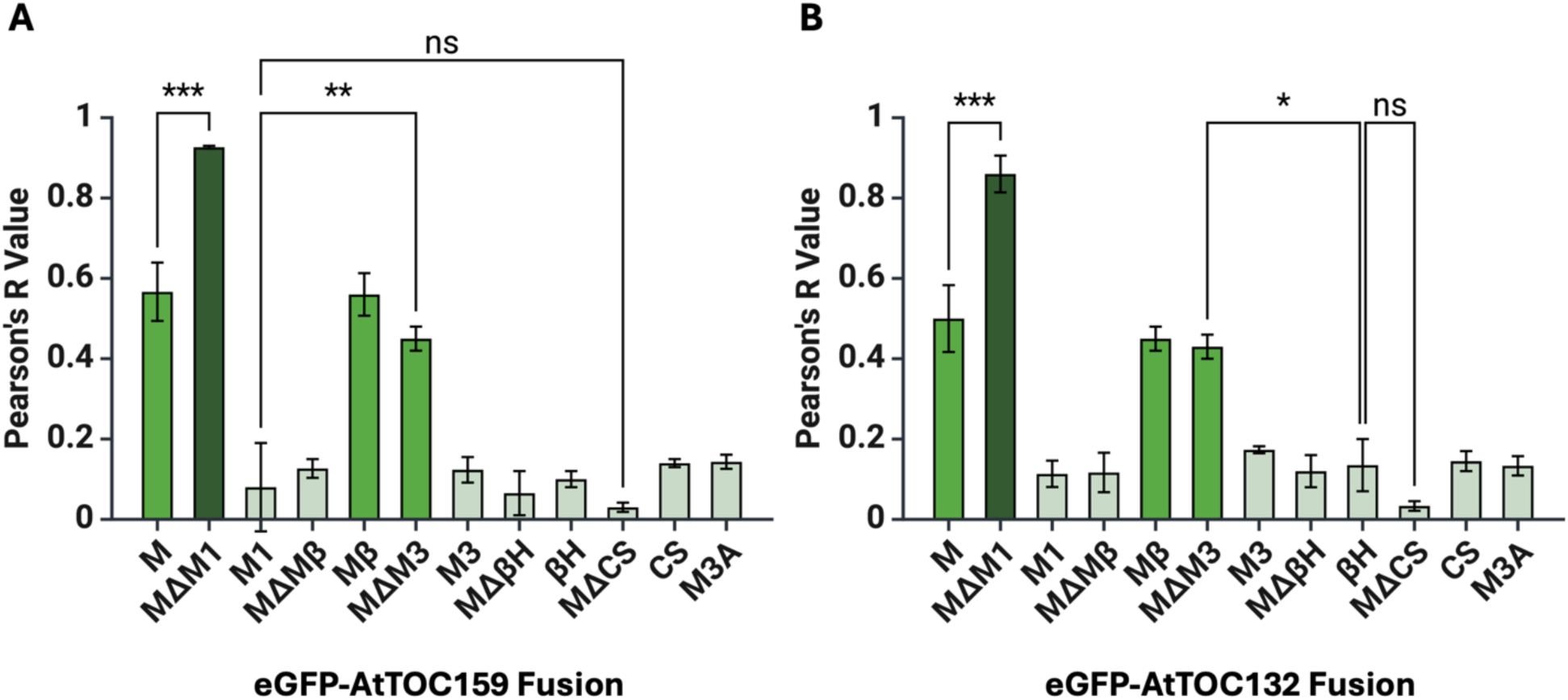
Plastid Colocalization Analysis of Transiently Expressed eGFP-AtTOC159 and eGFP-AtTOC132 Fusion Constructs in *Allium cepa* (onion) Epidermal Cells. The region of interest (ROI) depicted in **Figures 1, 5, and S1** were used in “Coloc 2” analysis with Image J (Fiji) software to determine the Pearson’s R value or correlation between eGFP and DsRed2 signals in merged fluorescent images for eGFP-AtTOC159 (**A**) and eGFP-AtTOC132 (**B**) fusion constructs (Schindelin et al., 2012). The Pearson’s R values depicted represent the mean (n = 3) ± the standard error of the mean (SEM). Statistical analyses using one-way ANOVA and the Tukey multiple comparisons test with the BioRender R 4.2.2 Graph integration demonstrate the same 3 significant populations for both eGFP-AtTOC159 and eGFP-AtTOC132 fusion constructs. One asterisk (*) represents a p-value ≤ 0.05, two asterisks (**) represents a p-value ≤ 0.01, three asterisks (***) represents a p-value ≤ 0.001, four asterisks (****) represents a p-value ≤ 0.0001 and ns represents a p-value > 0.05. Constructs showed either good plastid targeting capability (dark green), reduced targeting capability (green) or poor targeting capability (light green).

Since *A. cepa* epidermal cells lack true chloroplasts, containing etioplasts instead (Köhler and Hanson, 2000), chloroplast outer envelope targeting of key eGFP fusion constructs was confirmed in *A. thaliana* protoplasts. The same trends were observed, as seen in **Figure 6**, where protoplasts transformed with the AtTOC159MΔM1 construct showed tight rings of eGFP signal wrapped around the chloroplasts with little to no cytosolic eGFP signal. Protoplasts transformed with the AtTOC159MΔM3 construct which lacks the M3 subdomain resulted in reduced targeting capability, where eGFP signal was still associated with the chloroplasts, but a large amount of cytosolic eGFP signal between chloroplasts was also observed. Further, truncation of the Mβ subdomain (AtTOC159MΔMβ) and disruption of the β-signal (AtTOC159M3A) completely abolished chloroplast outer envelope targeting as protoplasts transformed with these constructs showed no eGFP signal associated with the chloroplasts. Additionally, the AtTOC159M3A construct showed irregularly shaped puncta of eGFP signal, which is likely a result of the inability of AtTOC159M3A to be properly targeted to and inserted in the chloroplast outer membrane, resulting in aggregation. These results were also observed for AtTOC132 eGFP fusion constructs (**Figure S3**).

**Figure 6.**
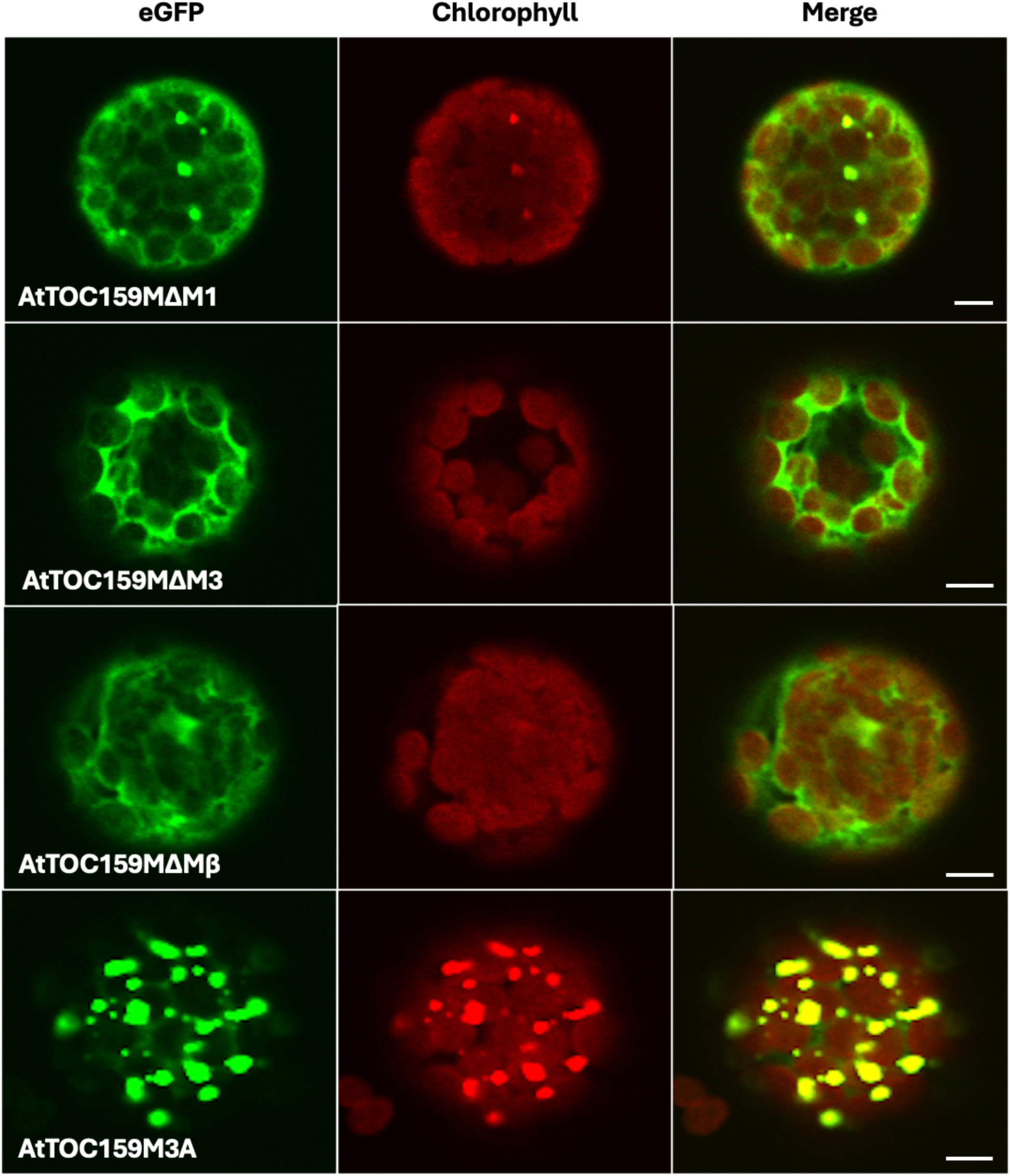
Subcellular Localization of Transiently Expressed eGFP-AtTOC159 Fusion Constructs Containing the M Domain, Subdomains and Subdomain Truncations in *Arabidopsis thaliana* Protoplasts. *Arabidopsis thaliana* protoplasts were isolated and transfected with eGFP-AtTOC159 fusion constructs (as depicted and described in **Figures 1A** and **2A**) using polyethylene glycol (PEG)-mediated transfection. Representative images for each transfection include eGFP (green), chlorophyll autofluorescence (red) and the merged channels. Scale bar = 10 µm.

### 3.3 AtTOC159 Receptor Targeting is Mediated by Interactions with Galactolipids

To better understand the mechanism by which the M3 subdomain imparts chloroplast outer envelope targeting specificity and considering the amphipathic and hydrophobic nature of the predicted ⍺-helix in this subdomain, we hypothesized that interactions with galactolipids unique to the chloroplast outer membrane could serve as a recognition site for the M3 subdomain. Interactions between the M3 subdomain and galactolipid-containing membranes were analyzed using CD spectroscopy and surface tension measurements of lipid monolayers deposited on a Langmuir-Blodgett trough. The AtTOC159M3 peptide was chemically synthesized, and its secondary structure observed in buffer and the presence of different lipid membranes. **Figure 7A** shows that AtTOC159M3 was disordered in buffer with a characteristic minimum below 200 nm (Greenfield, 2006; Miles et al., 2021). In the presence of POPC liposomes, AtTOC159M3 remained largely disordered, with a CD spectrum similar to that in buffer, but the appearance of a second, shallow minima around 222 nm was observed indicating some structural changes. In the presence of liposomes with a lipid composition that mimics the chloroplast outer envelope, AtTOC159M3 adopted an ⍺- helical structure, with characteristic minima at 208 nm and 222 nm (Greenfield, 2006; Miles et al., 2021). The role of each individual galactolipid in inducing the ⍺-helical conformation of AtTOC159M3 was also examined (**Figure 7B**). AtTOC159M3 adopted an ⍺-helical structure in the presence of POPC liposomes containing each of the galactolipids individually, although mean residue ellipticity was enhanced in those liposomes which contained SQDG.

**Figure 7.**
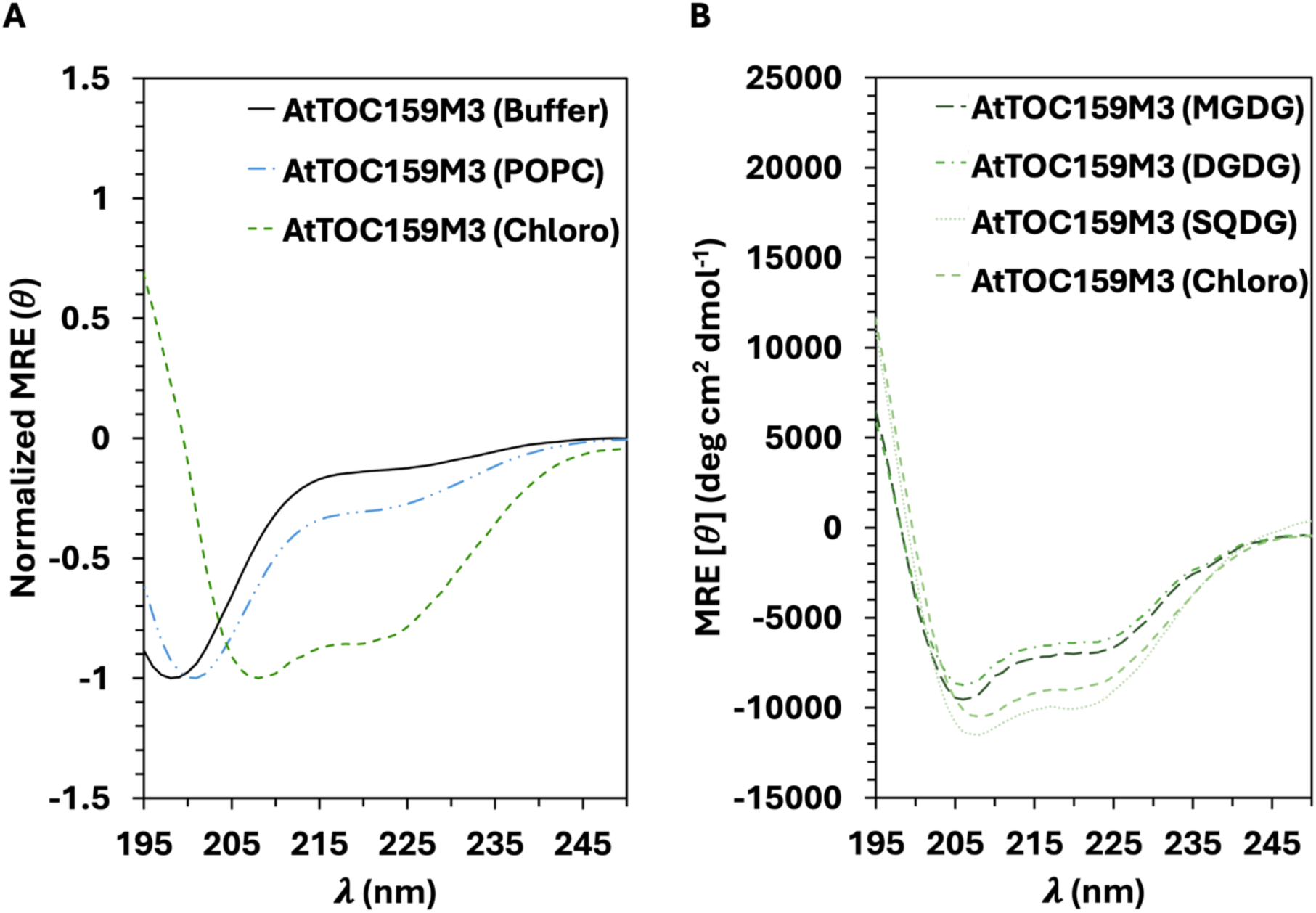
Structural Characterization of the C-Terminal Transit Peptide-Like Sequence of AtTOC159 in Dicerent Lipid Environments. (**A**) Far-UV CD spectra of the synthetic AtTOC159M3 peptide in Tris-NaF buqer (solid black line), POPC liposomes (blue dotted-dashed line) and liposomes which mimic the composition of the chloroplast outer envelope (Chloro, green dashed line). They y-axis is normalized for comparison of spectral changes. The spectra demonstrate a transition from random structure in buqer, partial structure in the presence of POPC liposomes and ⍺-helical structure in the presence of liposomes which mimic the chloroplast outer envelope. (**B**) Far-UV CD spectra of the synthetic AtTOC159M3 peptide in the presence of POPC liposomes containing 5% MGDG (long dashed line), 5% DGDG (dotted-dashed line), 5% SQDG (short dashed line) and liposomes which mimic the composition of the chloroplast outer envelope (Chloro, dotted line). All spectra indicate ⍺-helical structure, where the presence of SQDG increased helicity. The peptide concentration in (**A-B**) was 10 µM. All spectra are an average of 9 scans (n = 3) and were collected at 22 ℃ in a quartz cuvette with a 0.1 cm pathlength. MRE is the mean residue ellipticity (*θ*) in degrees cm^2^ dmol^-1^.

Lipid monolayers composed of different lipids were deposited on a Langmuir Blodgett trough and compressed at increasing initial pressures. As the AtTOC159M3 peptide was added to the subphase, interactions between the peptide and the lipid monolayer caused changes in surface tension (**Figure 8**). The maximum insertion pressure represents the initial pressure of the lipid monolayer at which the peptide can no longer interact with the lipid monolayer to cause observable changes in surface tension. The maximum insertion pressure is theref-ore a reflection of the affinity of the peptide for that lipid monolayer (Calvez et al., 2011; Sugár et al., 2005). To compare the affinity of the AtTOC159M3 peptide for specific lipid components, the maximum insertion pressure was determined for monolayers consisting of POPC or each galactolipid individually, and for a combination of lipids meant to mimic the composition of the chloroplast outer membrane (**Figure 8 inset**). The maximum insertion pressure for POPC monolayers was 15.608 mN/m. The maximum insertion pressure for MGDG monolayers, DGDG monolayers and monolayers that mimic the lipid composition of the chloroplast outer envelope were significantly higher than POPC monolayers at 20.713 mN/m, 20.944 mN/m and 22.498 mN/m, respectively. The maximum insertion pressure for SQDG monolayers was significantly higher than MGDG, DGDG and chloroplast outer envelope monolayers at 24.831 mN/m. Taken together, the CD spectroscopy and Langmuir Blodgett trough data suggest that the M3 subdomain participates in preferential interactions with galactolipid-containing membranes that are stabilized by the ⍺-helical conformation.

**Figure 8.**
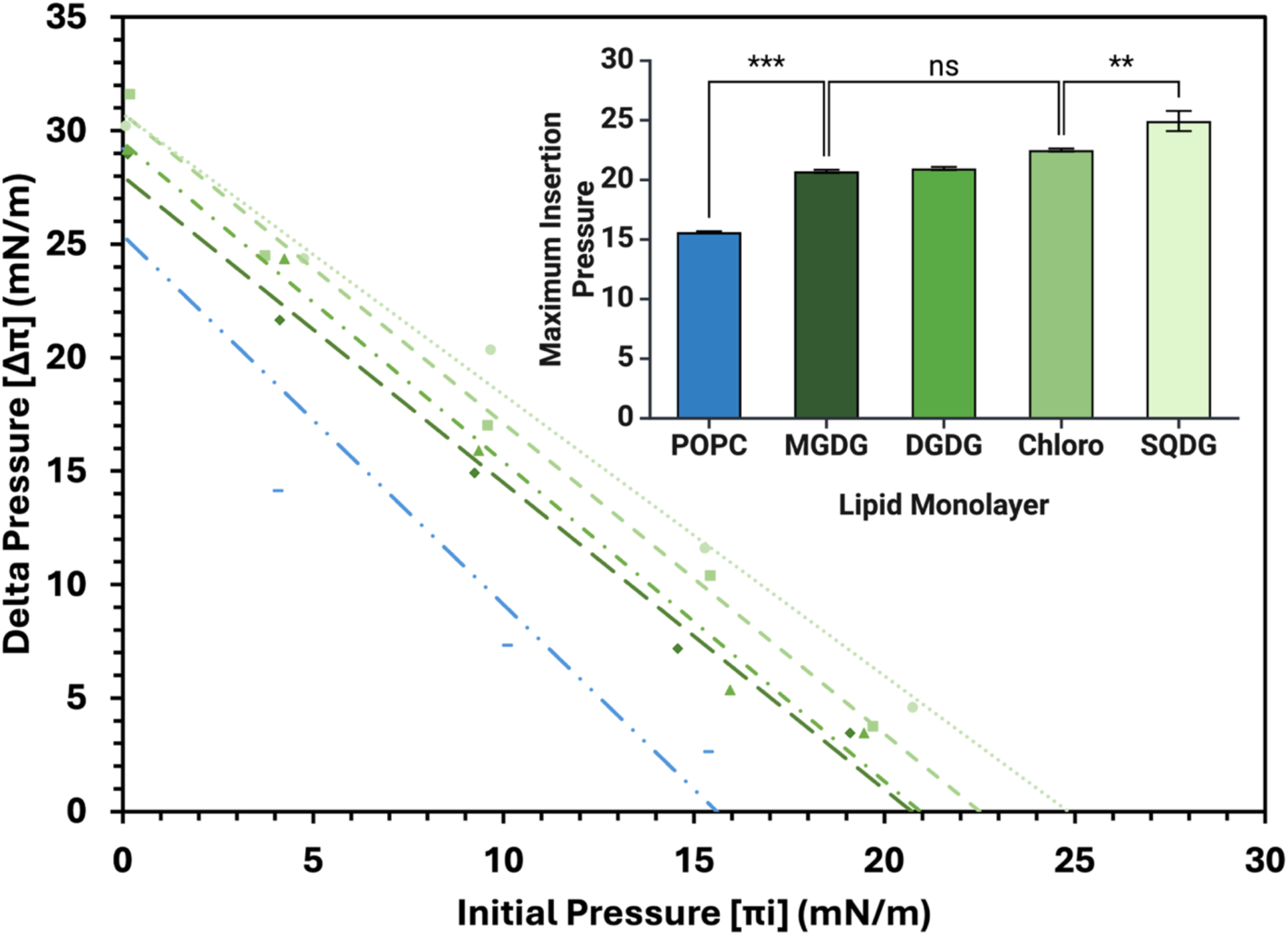
Maximum Insertion Pressure of the C-Terminal Transit Peptide-Like Sequence of AtTOC159 in Dicerent Lipid Environments. Lipid monolayers of POPC (blue double dotted dashed line), MGDG (green long dashed line), DGDG (green dot dashed line), SQDG (green short dashed line) and a lipid composition which mimics the chloroplast outer envelope (Chloro, green dotted line) were deposited on a Langmuir-Blodgett trough and changes in surface tension were measured in the presence of the synthetic AtTOC159M3 peptide at increasing initial pressures (**Figure S4**). Delta pressure was plotted against initial pressure for each lipid system and maximum insertion pressure (inset) was determined by extrapolation of the line of best fit to the x-intercept. Statistical analyses using one-way ANOVA and the Tukey multiple comparisons test with the BioRender R 4.2.2 Graph integration demonstrate significant increases in maximum insertion pressure that suggest the AtTOC159M3 peptide has a higher aqinity for galactolipids in the chloroplast outer envelope compared to POPC, with the most aqinity for SQDG. Two asterisks (**) represents a p-value ≤ 0.01, three asterisks (***) represents a p-value ≤0.001, and ns represents a p-value > 0.05. The peptide concentration was 10 µM and measurements were collected at 22 ℃. Maximum insertion pressure (inset) represents the mean (n = 3) ± the standard error of the mean (SEM). Delta pressure (Δπ), initial pressure (πi) and maximum insertion pressure are in mN/m.

## 4. Discussion

The results of this study contribute to our understanding of the complex targeting mechanism employed by AtTOC159 receptors (**Figure 9**), expanding our understanding of chloroplast outer membrane protein targeting and integration more generally. First, the structural prediction of AtTOC159, particularly the M domain, confirms the presence of a β- barrel that anchors the receptor in the chloroplast outer membrane. This structural feature, consistent across the AtTOC family, aligns with previous findings in *C. reinhardtii* and suggests an evolutionarily conserved mechanism for receptor integration. The fusion of this β-barrel with that of AtTOC75 introduces a novel hybrid channel concept, providing further evidence of functional interplay between these integral proteins. Such integration could imply that AtTOC159 has a more active role in preprotein translocation than previously thought, which warrants further investigation into its role within the import pore.

**Figure 9.**
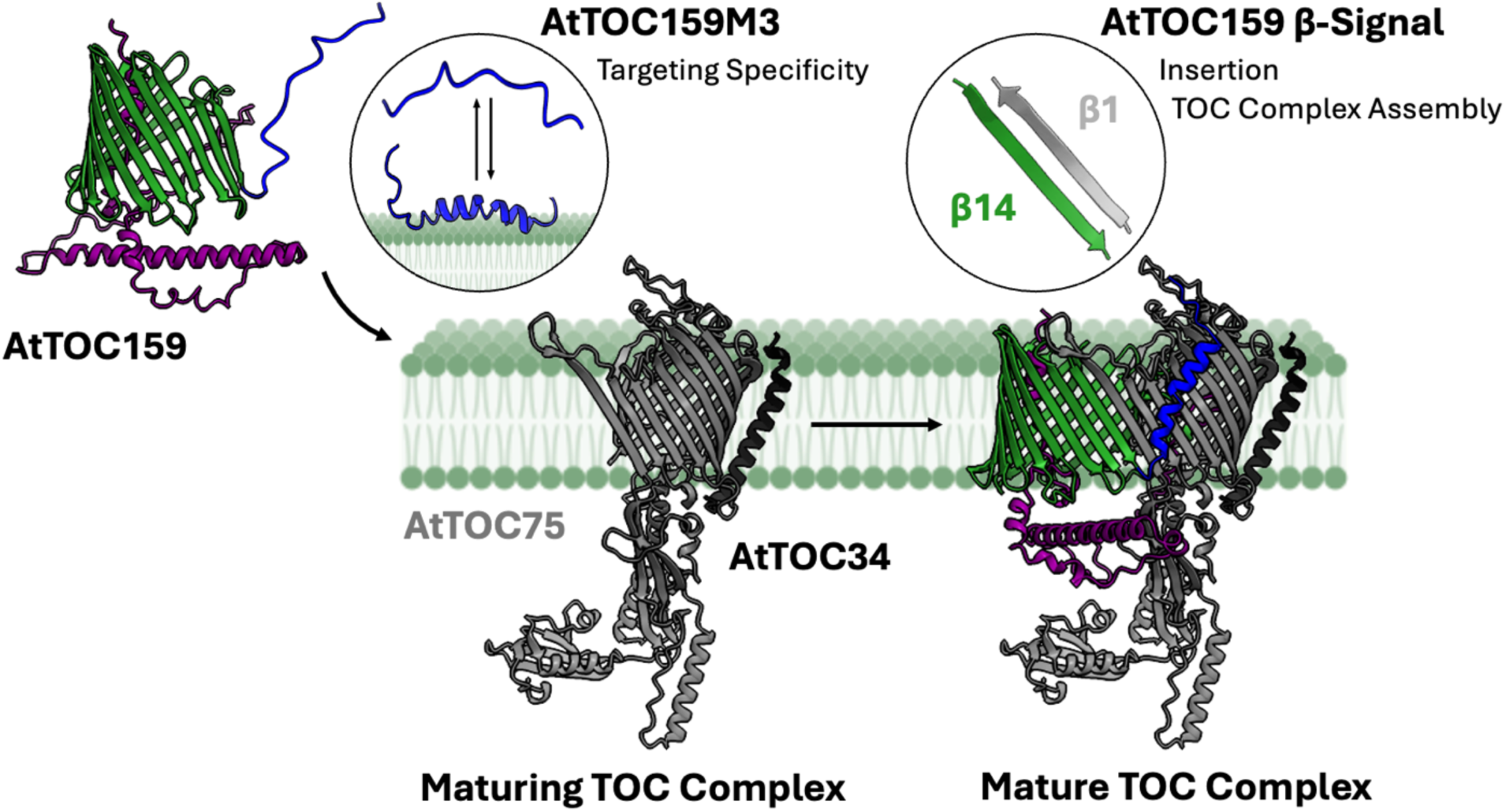
Proposed Model for Targeting of AtTOC159 Receptors to the Chloroplast Outer Envelope and Integration into Maturing TOC Complexes. After AtTOC159 receptors are translated in the cytosol, the disordered transit peptide-like sequence at the C-terminus (AtTOC159M3) interacts preferentially with galactolipids in the chloroplast outer membrane by stabilizing the formation of the membrane-interacting ⍺-helix. As AtTOC159 receptors encounter maturing TOC complexes, the β-signal drives their insertion into the chloroplast outer membrane and integration into the TOC complex. Protein structures were generated using UCSF Chimera.

A key finding in this study is the identification of a bi-partite targeting signal at the C-terminus of AtTOC159 and AtTOC132, composed of a β-signal and a TP-like sequence. Mutational analyses and truncation experiments demonstrated that the β-signal is critical for chloroplast outer membrane targeting, as its disruption abolishes localization. The conserved β-strand sequence (G-Q-Φ-[ST]-Φ-[RK]-X-[SN]-[ST]) across plant species suggests a targeting pathway similar to that observed in the insertion of β-barrel proteins in mitochondria and gram-negative bacteria. In contrast, the TP-like sequence enhances but is not necessary or sufficient for targeting, supporting its role as an auxiliary signal. This finding mirrors the behavior of TPs in chloroplast preprotein import, which often rely on multiple sub-domains for efficient targeting and have demonstrated specific interactions with MGDG, SQDG and PG (Bruce, 2000).

The role of chloroplast membrane-specific galactolipids in receptor targeting was another significant discovery. CD spectroscopy and Langmuir-Blodgett trough experiments revealed that the TP-like element interacts preferentially with lipids enriched in the chloroplast outer membrane, particularly SQDG, enhancing the α-helical structure of the peptide. The higher affinity of the TP-like sequence for SQDG compared to other lipids could be explained by the negatively charged lipid head group, considering the positively charged hydrophilic face of the AtTOC159M3 ⍺-helix. These lipid interactions may reflect a broader mechanism by which the unique lipid composition of the chloroplast membrane aids in the recognition and stabilization of other chloroplast outer membrane targeted proteins, providing an additional layer of selectivity in membrane protein targeting within the complex environment of the plant cell, which contains multiple organelles of endosymbiotic origin. The role of galactolipids, particularly in facilitating protein targeting, opens new avenues for exploring how membrane composition influences protein import and organelle biogenesis in photosynthetic eukaryotes.

While this study elucidates the targeting and membrane integration of TOC159 receptors in plants, several questions remain regarding their interaction with the maturing TOC complex and its broader role in chloroplast biogenesis. Future research should focus on the detailed structural dynamics of the TOC159 and TOC75 interaction. The hybrid β-barrel channel formed by these two proteins likely plays a critical role in preprotein import, but how these two β-barrels cooperate during the translocation process remains unclear. Cryo-electron microscopy could help resolve the structure of the TOC complex in plants in more detail, providing insights into how TOC159 receptors and TOC75 function together to facilitate preprotein translocation. Additionally, it would be beneficial to identify the cytosolic factors that recognize the β-signal and TP-like sequence of TOC159 receptors to direct them to the chloroplast outer membrane. Identifying these factors would clarify the steps involved in TOC159 receptor localization and could reveal parallels to mitochondrial and bacterial β- barrel insertion systems, such as the Sam50 and BamA complexes. Finally, this study raises important questions about whether similar targeting mechanisms are used by other chloroplast outer envelope proteins. Investigating the targeting signals of these proteins and their interactions with the chloroplast membrane would provide a more comprehensive understanding of how the entire outer envelope (proteome) is assembled and maintained. Additionally, it would be informative to explore whether other classes of outer envelope proteins utilize lipid-mediated targeting strategies similar to those observed for TOC159 receptors. Together, these lines of inquiry would significantly advance our understanding of chloroplast membrane protein targeting and its broader implications for chloroplast function and plant biology.

## 6. Author Contributions

MF: Conceptualization, Formal Analysis, Investigation, Methodology, Validation, Visualization, Writing – original draft, Writing – review & editing; GS: Formal Analysis, Investigation, Methodology, Validation, Writing – review & editing; SDXC: Funding Acquisition, Resources, Supervision, Writing – review & editing; MJN: Funding Acquisition, Resources, Supervision, Writing – review & editing; MDS: Funding Acquisition, Resources, Supervision, Writing – review & editing.

## 8. Acknowledgements

We would like to thank Mohsen Shojaei Barjouei (Undergraduate Volunteer, Wilfrid Laurier University) for his care and consistency in *A. thaliana* planting and maintenance; Max Pottier (Instrumentation Technician, Faculty of Science, Wilfrid Laurier University) for his technical assistance operating the confocal fluorescence microscope and other instruments; Justin Hume (Undergraduate Thesis Student and Research Assistant, Wilfrid Laurier University) and Noah Gilholm (NSERC USRA, Wilfrid Laurier University) for their work in preparing plasmid DNA required for protoplast transfection experiments.

