## Supplemental Material for "AtTOC159 Receptors are Targeted to the Chloroplast Outer Membrane by a β-Signal and Galactolipid-Specific Transit Peptide-Like Sequence at the C-Terminus"

### Supplementary Material

**Table 1. eGFP-AtTOC159 Construct Sequences.** Amino acid residue sequences from AtTOC159 fused to the C-terminus of eGFP are included. The residues within the β-signal consensus sequence of AtTOC159M3A that have been mutated to alanine (A) are bolded.

| eGFP Construct | Amino Acid Sequence |
|---|---|
| AtTOC159M | VRSPPLPYLLSWLLQSRAHPKLPGDQGGDSVDSDIEIDDVSDSEQEDGEDDEYDQLPPFKPLRKTQLAKLSNEQR<br>KAYFEEYDYRVKLLQKKQWREELKRMKEMKKNGKKLGESEFGYPGEEDDPENGAPAAVPVPLPDMVLPPSFDSD<br>NSAYRYRYLEPTSQLLTRPVLDTHGWDHDCGYDGVNAEHSLALASRFPATATVQVTKDKKEFNIHLDSSVSAKHGE<br>NGSTMAGFDIQNVGKQLAYVVRGETKFKNLRKNKTTVGGSVTFLGENIATGVKLEDQIALGKRLVLVGSTGTMRSQ<br>GDSAYGANLEVRLREADFPIGQDQSSFGLSLVKWRGDLALGANLQSQVSVGRNSKIALRAGLNNKMS**G**QITV**R**TS<br>**S**SDQLQIALTAILPIAMSIYKSIRPEATNDKYSMY |
| AtTOC159M1 | VRSPPLPYLLSWLLQSRAHPKLPGDQGGDSVDSDIEIDDVSDSEQEDGEDDEYDQLPPFKPLRKTQLAKLSNEQR<br>KAYFEEYDYRVKLLQKKQWREELKRMKEMKKNGKKLGESEFGYPGEEDDPENGAPAAVPVPLPDMVLPPSFDSD<br>NS |
| AtTOC159Mβ | AYRYRYLEPTSQLLTRPVLDTHGWDHDCGYDGVNAEHSLALASRFPATATVQVTKDKKEFNIHLDSSVSAKHGEN<br>GSTMAGFDIQNVGKQLAYVVRGETKFKNLRKNKTTVGGSVTFLGENIATGVKLEDQIALGKRLVLVGSTGTMRSQG<br>DSAYGANLEVRLREADFPIGQDQSSFGLSLVKWRGDLALGANLQSQVSVGRNSKIALRAGLNNKMS**G**QITV**R**TS**S** |
| AtTOC159M3 | SDQLQIALTAILPIAMSIYKSIRPEATNDKYSMY |
| AtTOC159MΔM1 | AYRYRYLEPTSQLLTRPVLDTHGWDHDCGYDGVNAEHSLALASRFPATATVQVTKDKKEFNIHLDSSVSAKHGEN<br>GSTMAGFDIQNVGKQLAYVVRGETKFKNLRKNKTTVGGSVTFLGENIATGVKLEDQIALGKRLVLVGSTGTMRSQG<br>DSAYGANLEVRLREADFPIGQDQSSFGLSLVKWRGDLALGANLQSQVSVGRNSKIALRAGLNNKMS**G**QITV**R**TS**S**<br>SDQLQIALTAILPIAMSIYKSIRPEATNDKYSMY |
| AtTOC159MΔMβ | RSPPLPYLLSWLLQSRAHPKLPGDQGGDSVDSDIEIDDVSDSEQEDGEDDEYDQLPPFKPLRKTQLAKLSNEQRK<br>AYFEEYDYRVKLLQKKQWREELKRMKEMKKNGKKLGESEFGYPGEEDDPENGAPAAVPVPLPDMVLPPSFDSDN<br>SSDQLQIALTAILPIAMSIYKSIRPEATNDKYSMY |
| AtTOC159MΔM3 | VRSPPLPYLLSWLLQSRAHPKLPGDQGGDSVDSDIEIDDVSDSEQEDGEDDEYDQLPPFKPLRKTQLAKLSNEQR<br>KAYFEEYDYRVKLLQKKQWREELKRMKEMKKNGKKLGESEFGYPGEEDDPENGAPAAVPVPLPDMVLPPSFDSD<br>NSAYRYRYLEPTSQLLTRPVLDTHGWDHDCGYDGVNAEHSLALASRFPATATVQVTKDKKEFNIHLDSSVSAKHGE<br>NGSTMAGFDIQNVGKQLAYVVRGETKFKNLRKNKTTVGGSVTFLGENIATGVKLEDQIALGKRLVLVGSTGTMRSQ<br>GDSAYGANLEVRLREADFPIGQDQSSFGLSLVKWRGDLALGANLQSQVSVGRNSKIALRAGLNNKMS**G**QITV**R**TS<br>**S** |
| AtTOC159βH | KIALRAGLNNKMS**G**QITV**R**TS**S** |
| AtTOC159MΔβH | VRSPPLPYLLSWLLQSRAHPKLPGDQGGDSVDSDIEIDDVSDSEQEDGEDDEYDQLPPFKPLRKTQLAKLSNEQR<br>KAYFEEYDYRVKLLQKKQWREELKRMKEMKKNGKKLGESEFGYPGEEDDPENGAPAAVPVPLPDMVLPPSFDSD<br>NSAYRYRYLEPTSQLLTRPVLDTHGWDHDCGYDGVNAEHSLALASRFPATATVQVTKDKKEFNIHLDSSVSAKHGE<br>NGSTMAGFDIQNVGKQLAYVVRGETKFKNLRKNKTTVGGSVTFLGENIATGVKLEDQIALGKRLVLVGSTGTMRSQ<br>GDSAYGANLEVRLREADFPIGQDQSSFGLSLVKWRGDLALGANLQSQVSVGRNSSDQLQIALTAILPIAMSIYKSIR<br>PEATNDKYSMY |
| AtTOC159CS | MS**G**QITV**R**TS**S**SDQLQIALT |
| AtTOC159MΔCS | VRSPPLPYLLSWLLQSRAHPKLPGDQGGDSVDSDIEIDDVSDSEQEDGEDDEYDQLPPFKPLRKTQLAKLSNEQR<br>KAYFEEYDYRVKLLQKKQWREELKRMKEMKKNGKKLGESEFGYPGEEDDPENGAPAAVPVPLPDMVLPPSFDSD<br>NSAYRYRYLEPTSQLLTRPVLDTHGWDHDCGYDGVNAEHSLALASRFPATATVQVTKDKKEFNIHLDSSVSAKHGE<br>NGSTMAGFDIQNVGKQLAYVVRGETKFKNLRKNKTTVGGSVTFLGENIATGVKLEDQIALGKRLVLVGSTGTMRSQ<br>GDSAYGANLEVRLREADFPIGQDQSSFGLSLVKWRGDLALGANLQSQVSVGRNSKIALRAGLNNKAILPIAMSIYK<br>SIRPEATNDKYSMY |
| AtTOC159M3A | VRSPPLPYLLSWLLQSRAHPKLPGDQGGDSVDSDIEIDDVSDSEQEDGEDDEYDQLPPFKPLRKTQLAKLSNEQR<br>KAYFEEYDYRVKLLQKKQWREELKRMKEMKKNGKKLGESEFGYPGEEDDPENGAPAAVPVPLPDMVLPPSFDSD<br>NSAYRYRYLEPTSQLLTRPVLDTHGWDHDCGYDGVNAEHSLALASRFPATATVQVTKDKKEFNIHLDSSVSAKHGE<br>NGSTMAGFDIQNVGKQLAYVVRGETKFKNLRKNKTTVGGSVTFLGENIATGVKLEDQIALGKRLVLVGSTGTMRSQ<br>GDSAYGANLEVRLREADFPIGQDQSSFGLSLVKWRGDLALGANLQSQVSVGRNSKIALRAGLNNKMS**A**QITV**A**TS<br>**A**SDQLQIALTAILPIAMSIYKSIRPEATNDKYSMY |

**Table 2. eGFP-AtTOC132 Construct Sequences.** Amino acid residue sequences from AtTOC132 fused to the C-terminus of eGFP are included. The residues within the β-signal consensus sequence of AtTOC132M3A that have been mutated to alanine (A) are bolded.

| eGFP Construct | Amino Acid Sequence |
|---|---|
| AtTOC132M | DNIPGRPFAARSKAPPLPFLLSSLLQSRPQPKLPEQQYGDEEDEDDLEESSDSDEESEYDQLPPFKSLTKAQMATL<br>SKSQKKQYLDEMEYREKLLMKKQMKEERKRRKMFKKFAAEIKDLPDGYSENVEEESGGPASVPVPMPDLSLPASF<br>DSDNPTHRYRYLDSSNQWLVRPVLETHGWDHDIGYEGVNAERLFVVKEKIPISVSGQVTKDKKDANVQLEMASS<br>VKHGEGKSTSLGFDMQTVGKELAYTLRSETRFNNFRRNKAAAGLSVTHLGDSVSAGLKVEDKFIASKWFRIVMSG<br>GAMTSRGDFAYGGTLEAQLRDKDYPLGRFLTTLGLSVMDWHGDLAIGGNIQSQVPIGRSSNLIARANLNNRGA**G**<br>QVSV**R**VN**S**SEQLQLAMVAIVPLFKKLLSYYYPQTQYGQ |
| AtTOC132M1 | DNIPGRPFAARSKAPPLPFLLSSLLQSRPQPKLPEQQYGDEEDEDDLEESSDSDEESEYDQLPPFKSLTKAQMATL<br>SKSQKKQYLDEMEYREKLLMKKQMKEERKRRKMFKKFAAEIKDLPDGYSENVEEESGGPASVPVPMPDLSLPASF<br>DSDNP |
| AtTOC132Mβ | THRYRYLDSSNQWLVRPVLETHGWDHDIGYEGVNAERLFVVKEKIPISVSGQVTKDKKDANVQLEMASSVKHGE<br>GKSTSLGFDMQTVGKELAYTLRSETRFNNFRRNKAAAGLSVTHLGDSVSAGLKVEDKFIASKWFRIVMSGGAMTSR<br>GDFAYGGTLEAQLRDKDYPLGRFLTTLGLSVMDWHGDLAIGGNIQSQVPIGRSSNLIARANLNNRGA**G**QVSV**R**V<br>N**S** |
| AtTOC132M3 | SEQLQLAMVAIVPLFKKLLSYYYPQTQYGQ |
| AtTOC132MΔM1 | THRYRYLDSSNQWLVRPVLETHGWDHDIGYEGVNAERLFVVKEKIPISVSGQVTKDKKDANVQLEMASSVKHGE<br>GKSTSLGFDMQTVGKELAYTLRSETRFNNFRRNKAAAGLSVTHLGDSVSAGLKVEDKFIASKWFRIVMSGGAMTSR<br>GDFAYGGTLEAQLRDKDYPLGRFLTTLGLSVMDWHGDLAIGGNIQSQVPIGRSSNLIARANLNNRGA**G**QVSV**R**V<br>N**S**SEQLQLAMVAIVPLFKKLLSYYYPQTQYGQ |
| AtTOC132MΔMβ | DNIPGRPFAARSKAPPLPFLLSSLLQSRPQPKLPEQQYGDEEDEDDLEESSDSDEESEYDQLPPFKSLTKAQMATL<br>SKSQKKQYLDEMEYREKLLMKKQMKEERKRRKMFKKFAAEIKDLPDGYSENVEEESGGPASVPVPMPDLSLPASF<br>DSDNPSEQLQLAMVAIVPLFKKLLSYYYPQTQYGQ |
| AtTOC132MΔM3 | DNIPGRPFAARSKAPPLPFLLSSLLQSRPQPKLPEQQYGDEEDEDDLEESSDSDEESEYDQLPPFKSLTKAQMATL<br>SKSQKKQYLDEMEYREKLLMKKQMKEERKRRKMFKKFAAEIKDLPDGYSENVEEESGGPASVPVPMPDLSLPASF<br>DSDNPTHRYRYLDSSNQWLVRPVLETHGWDHDIGYEGVNAERLFVVKEKIPISVSGQVTKDKKDANVQLEMASS<br>VKHGEGKSTSLGFDMQTVGKELAYTLRSETRFNNFRRNKAAAGLSVTHLGDSVSAGLKVEDKFIASKWFRIVMSG<br>GAMTSRGDFAYGGTLEAQLRDKDYPLGRFLTTLGLSVMDWHGDLAIGGNIQSQVPIGRSSNLIARANLNNRGA**G**<br>QVSV**R**VN**S** |
| AtTOC132βH | NLIARANLNNRGA**G**QVSV**R**VN**S** |
| AtTOC132MΔβH | DNIPGRPFAARSKAPPLPFLLSSLLQSRPQPKLPEQQYGDEEDEDDLEESSDSDEESEYDQLPPFKSLTKAQMATL<br>SKSQKKQYLDEMEYREKLLMKKQMKEERKRRKMFKKFAAEIKDLPDGYSENVEEESGGPASVPVPMPDLSLPASF<br>DSDNPTHRYRYLDSSNQWLVRPVLETHGWDHDIGYEGVNAERLFVVKEKIPISVSGQVTKDKKDANVQLEMASS<br>VKHGEGKSTSLGFDMQTVGKELAYTLRSETRFNNFRRNKAAAGLSVTHLGDSVSAGLKVEDKFIASKWFRIVMSG<br>GAMTSRGDFAYGGTLEAQLRDKDYPLGRFLTTLGLSVMDWHGDLAIGGNIQSQVPIGRSSSEQLQLAMVAIVPLF<br>KKLLSYYYPQTQYGQ |
| AtTOC132CS | GA**G**QVSV**R**VN**S**SEQLQLAMV |
| AtTOC132MΔCS | DNIPGRPFAARSKAPPLPFLLSSLLQSRPQPKLPEQQYGDEEDEDDLEESSDSDEESEYDQLPPFKSLTKAQMATL<br>SKSQKKQYLDEMEYREKLLMKKQMKEERKRRKMFKKFAAEIKDLPDGYSENVEEESGGPASVPVPMPDLSLPASF<br>DSDNPTHRYRYLDSSNQWLVRPVLETHGWDHDIGYEGVNAERLFVVKEKIPISVSGQVTKDKKDANVQLEMASS<br>VKHGEGKSTSLGFDMQTVGKELAYTLRSETRFNNFRRNKAAAGLSVTHLGDSVSAGLKVEDKFIASKWFRIVMSG<br>GAMTSRGDFAYGGTLEAQLRDKDYPLGRFLTTLGLSVMDWHGDLAIGGNIQSQVPIGRSSNLIARANLNNRAIVPL<br>FKKLLSYYYPQTQYGQ |
| AtTOC132M3A | DNIPGRPFAARSKAPPLPFLLSSLLQSRPQPKLPEQQYGDEEDEDDLEESSDSDEESEYDQLPPFKSLTKAQMATL<br>SKSQKKQYLDEMEYREKLLMKKQMKEERKRRKMFKKFAAEIKDLPDGYSENVEEESGGPASVPVPMPDLSLPASF<br>DSDNPTHRYRYLDSSNQWLVRPVLETHGWDHDIGYEGVNAERLFVVKEKIPISVSGQVTKDKKDANVQLEMASS<br>VKHGEGKSTSLGFDMQTVGKELAYTLRSETRFNNFRRNKAAAGLSVTHLGDSVSAGLKVEDKFIASKWFRIVMSG<br>GAMTSRGDFAYGGTLEAQLRDKDYPLGRFLTTLGLSVMDWHGDLAIGGNIQSQVPIGRSSNLIARANLNNRGA**A**<br>QVSV**A**VN**A**SEQLQLAMVAIVPLFKKLLSYYYPQTQYGQ |

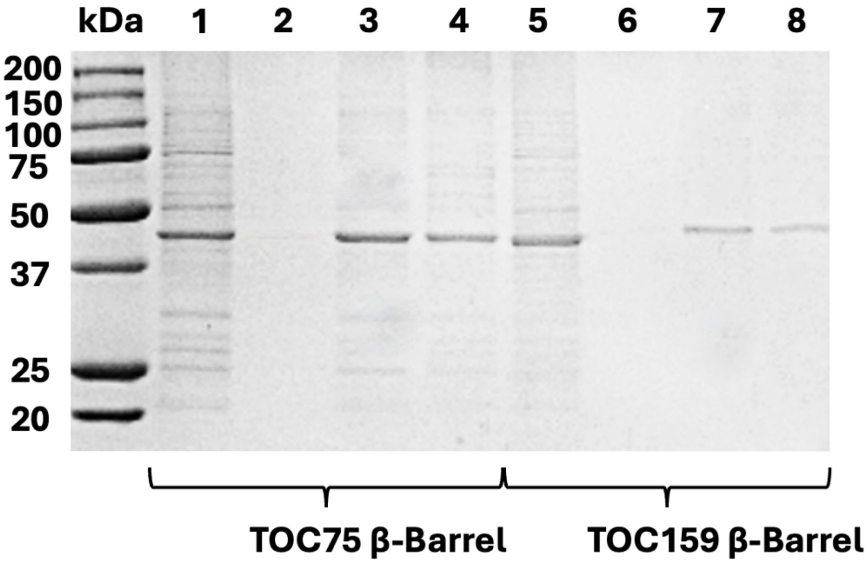


**Figure S1. SDS-PAGE Analysis of the Recombinant Expression and Purification of AtTOC75, AtTOC159 and AtTOC132 β-Barrels.** Recombinant proteins were expressed in *Escherichia coli* using IPTG induction and purified from inclusion bodies using immobilized metal affinity chromatography after on-column re-folding in 0.1% Triton X-100. (Lanes 1 and 5) Solubilized inclusion bodies. (Lanes 2 and 6) On-column re-folding wash. (Lanes 3 and 7) Elution from immobilized metal affinity chromatography column. (Lanes 4 and 8) Elution from size exclusion de-salt column. Proteins are separated by a 12% resolving gel and visualized by staining with Coomassie Brilliant Blue. The molecular weight ladder (kDa) used was the Precision Plus Protein™ All Blue Prestained Protein Standards from Bio-Rad (1610373).

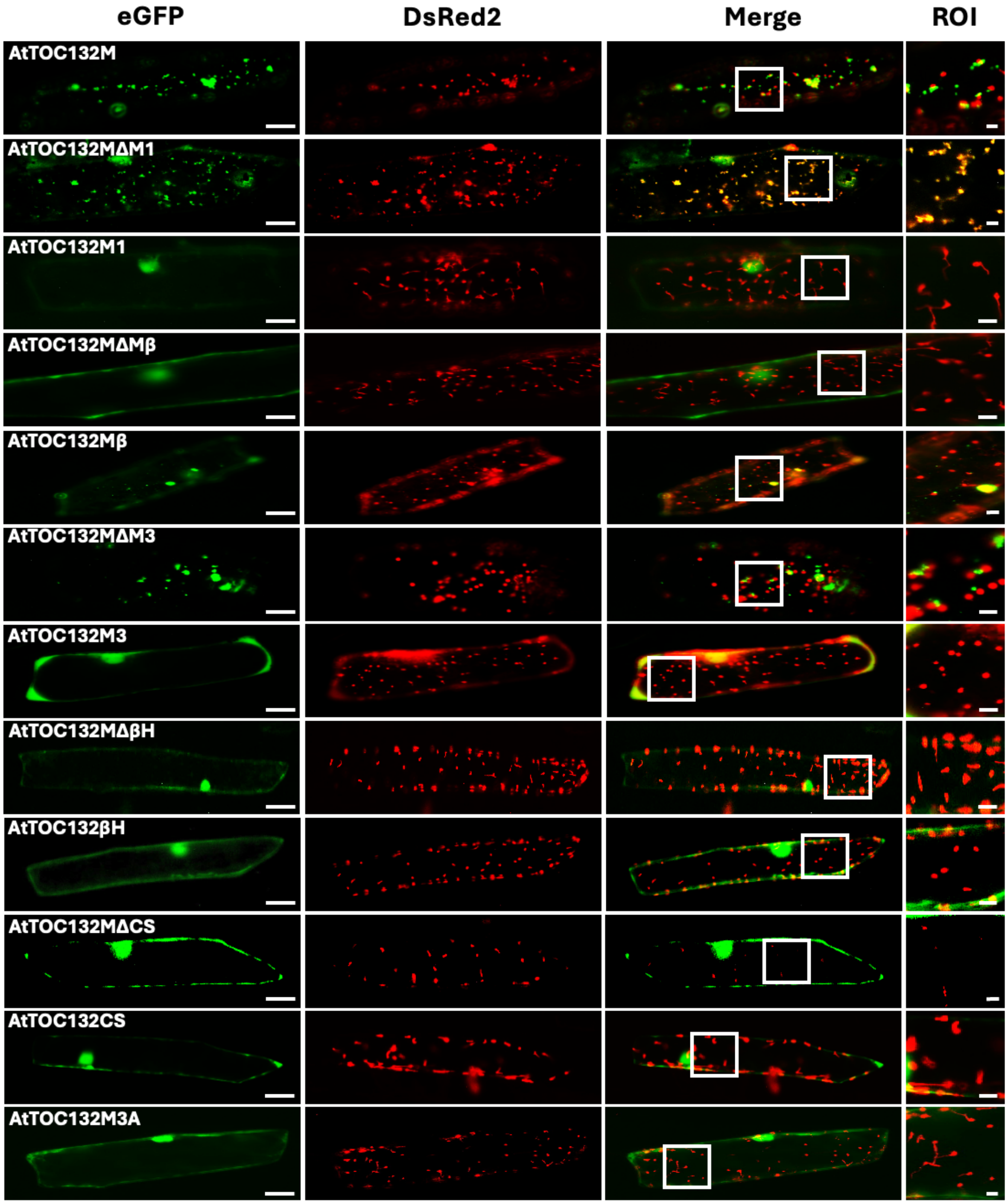


**Figure S2. Subcellular Localization of Transiently Expressed eGFP-AtTOC132 Fusion Constructs in *Allium cepa* (onion) Epidermal Cells.** AtTOC132 constructs were fused to the C-terminus of eGFP in the pSAT6-35S:eGFP-C1 vector to be transiently expressed using the 35S promoter. The epidermal cells of onion bulbs were co-transfected with eGFP-AtTOC132 fusion constructs and a DsRed2-Ferredoxin transit peptide fusion construct using biolistic bombardment. Representative images for each co-bombardment include eGFP (green), DsRed2 (red) and the merged channels. Colocalization of the green and red channels to plastids produce a yellow signal. The region of interest (ROI) is representative of the areas in the merged image used to analyze colocalization (**Figure 5**). Scale bars = 50 µm (eGFP) and 5 µm (ROI).

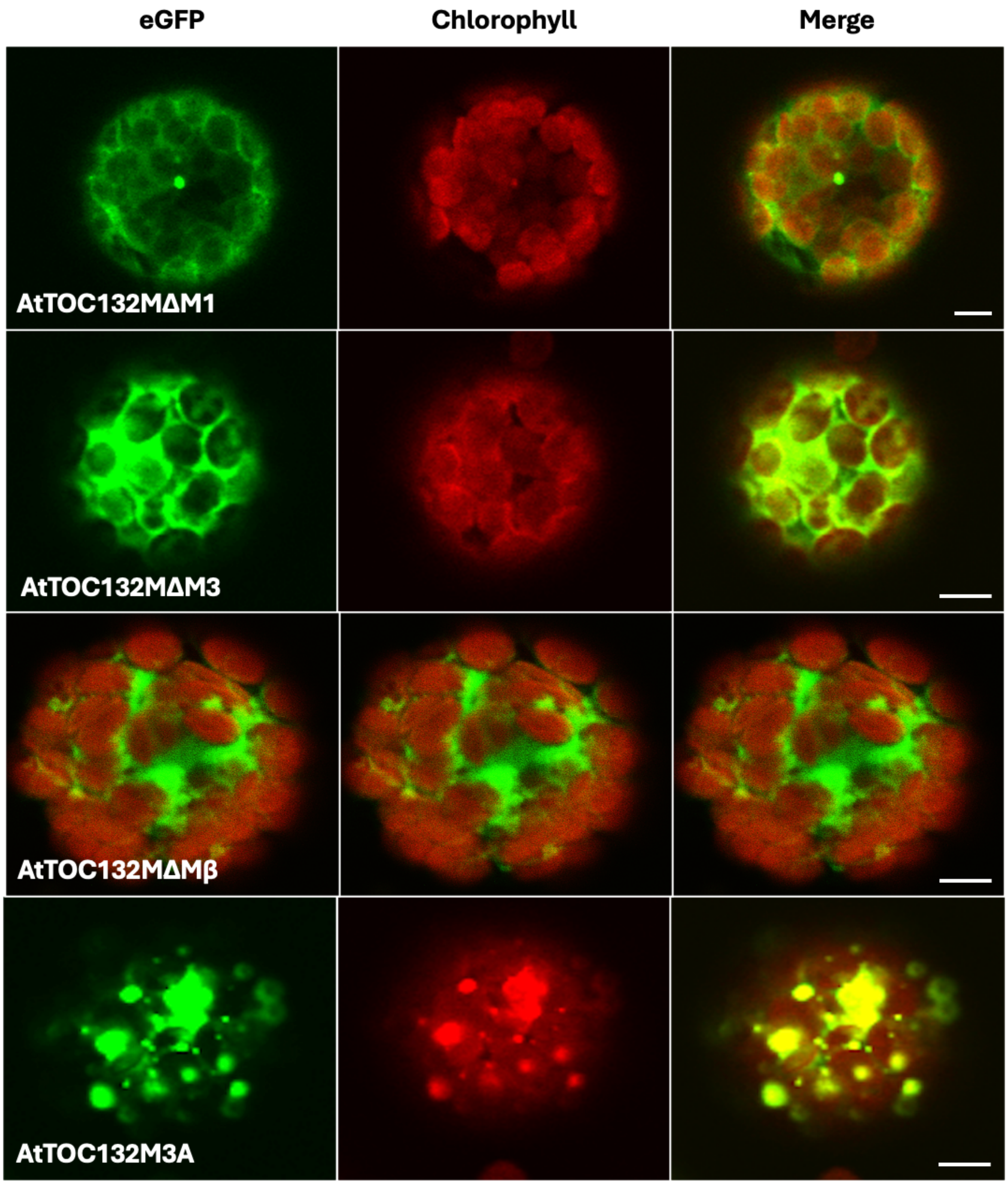


**Figure S3. Subcellular Localization of Transiently Expressed eGFP-AtTOC132 Fusion Constructs Containing the M Domain, Subdomains and Subdomain Truncations in *Arabidopsis thaliana* Protoplasts.** *Arabidopsis thaliana* protoplasts were isolated and transfected with eGFP-AtTOC132 fusion constructs (as depicted and described in **Figures 1A and 2A**) using polyethylene glycol (PEG)-mediated transfection. Representative images for each transfection include eGFP (green), chlorophyll autofluorescence (red) and the merged channels. Scale bar = 10 μm.

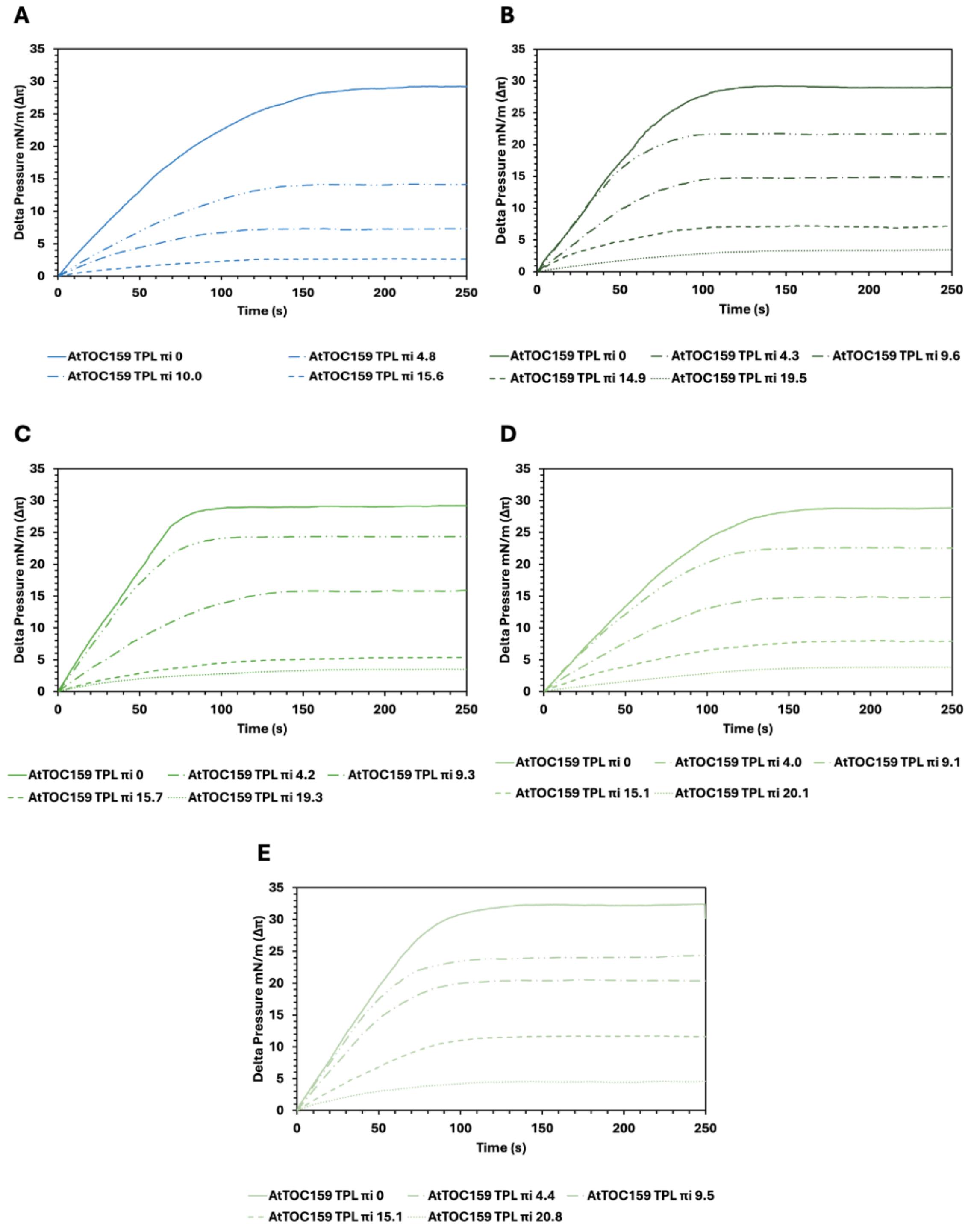


**Figure S4. Lipid Monolayer Equilibrium Pressure for AtTOC159M3 at Varying Initial Pressures.** Changes in surface tension of various lipid monolayers deposited on a Langmuir-Blodgett trough in the presence of the synthetic AtTOC159M3 peptide were measured as a function of time (s) at increasing initial pressures for POPC (A), MGDG (B), DGDG (C), a lipid composition which mimics the chloroplast outer envelope (D) and SQDG (E). Curves were generated from the mean of three delta pressure measurements at each initial pressure from separate experiments (n = 3). Delta pressure (Δπ) and initial pressure (πi) are in mN/m.
